# Coupled Nosé-Hoover equations of motions without time scaling

**DOI:** 10.1101/067397

**Authors:** Ikuo Fukuda, Kei Moritsugu

## Abstract

The Nosé-Hoover (NH) equation of motion is widely used in molecular dynamics simulations. It enables us to set a constant temperature and produce the canonical distribution for a target physical system. For the purpose of investigating the physical system under *fluctuating* temperature, we have introduced a *coupled Nosé-Hoover* equation in our previous work [*J. Phys. A* 48 455001 (2015)]. The coupled NH equation implements a fluctuating heat-bath temperature in the NH equation of the physical system, and also keeps a statistically complete description via an invariant measure of the total system composed of the physical system and a “temperature system”. However, a difficulty lies in that the time development of the physical system may not correspond to the realistic physical process, because of the need of a scaled time average to compute thermodynamical quantities. The current work gives a solution by presenting a new scheme, which is free from the scaled time but retains the statistical description. By use of simple model systems, we validate the current scheme and compare with the original scheme. The sampling property of the current scheme is also clarified to investigate the effect of function setting used for the distribution of the total system.

## I. INTRODUCTION

Molecular dynamics (MD) [1–3] is an efficient method to investigate the characteristics of physical systems in terms of both microscopic descriptions–based on atomic models–and their time developments–based on the Newtonian equations of motion (EOM). A realistic macroscopic system composed of many atoms and molecules can now be a target with a help of powerful computational architecture. However, the Newtonian EOM itself is not convenient for a direct comparison between MD simulations and experiments, because an experiment is done under the environment with a constant temperature (e.g., 300 K), while the Newtonian EOM does not provide the target temperature in general.

This problem is solved by the Nosé-Hoover (NH) equation [4, 5], which enables us to perform a constant temperature MD under a target temperature. The NH equation combines the Newtonian EOM with a heat-bath related friction variable, and realizes the equilibrium characterized by the canonical distribution at an arbitrary target temperature *T*_ex_. This is a physically-sound combination, and the results have been analyzed in various manners (see e.g., references cited in [6, 7]).

The NH equation is represented by

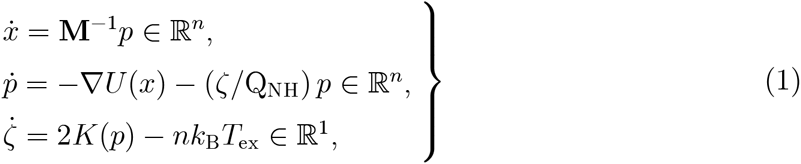

where *x* ≡ (*x*_1_, …, *x_n_*) ∈ ℝ^*n*^ are the coordinates for the physical system (PS) with *n* degrees of freedom, and *p* ≡ (*p*_1_, …, *p_n_*) ∈ ℝ^*n*^ are the corresponding momenta, *U*(*x*) is the potential energy describing an interaction among individual degrees of freedom, and *K*(*p*) is the kinetic energy. *ζ* ∈ ℝ is the heat-bath related friction variable to control the temperature of the PS and to set it to a given value *T*_ex_. *ζ* measures the difference between the instantaneous PS temperature 2*K*(*p*)/*nk*_B_ (*k*_B_ is Boltzmann’s constant) and the target temperature *T*_ex_, and the PS receives a feedback from the difference though the friction term −(*ζ*/Q_NH_)*p*. Here Q_NH_ is a positive parameter (called a Nosé’s mass), which plays a role of the friction coefficient and a role of coupling constant between the PS and the heat-bath [8]. The realization of the canonical distribution is due to the smooth measure 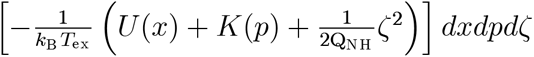, which is invariant for the EOM.

We often need to investigate the PS beyond the equilibrium or in nonequilibrium, such as heat flow [9] generated from two or more heat baths concerning with the Fourier law [10], and relaxation process after change of the system temperature [6, 11]. For the purpose of investigating the PS under non-constant temperature, *ad hoc* procedures for varying the temperature *T*_ex_ using the NH equation is possible. However, it provides no clear information about the probability distribution of the PS, which allows the comparison between the MD simulation and the experiments.

To solve this problem, we proposed the coupled NH (cNH) equation in our previous work [12, 13]. In the cNH equation, *T*_ex_ is replaced with a dynamical variable *T*(*x, p,𝒬*) that allows a fluctuation by coupling with another NH equation having its coordinate *𝒬*. This additional NH equation describes a *temperature system (TS)*, which consists of the *TS coordinate Q, the TS momentum 𝒫, and the TS friction variable η*. Namely, the cNH couples the NH equation for the PS and the NH equation for the TS. The interactions between these two systems are achieved through the *dynamical temperature T*(*x, p,𝒬*) governing the PS and through the *TS* potential energy *V_E_*(*x,p*)(*𝒬*) as a function of the TS degree of freedom *𝒬* as well as the PS energy *E*(*x,p*).

The cNH equation is not derived by an *ad hoc* manner but has a certain mathematical structure. In fact, the flow of the cNH has an invariant measure that is related to a density exp [−*β* (*U*(*x*) + *K*(*p*))] *f*(*β*) as a special case, where *β* is now a dynamical variable related to *𝒬*. This density indicates the relationship with the cNH and superstatistics [14–16]. Thus, we can fluctuate the heat-bath temperature of the PS and also obtain the statistical information of the PS. This is a solution to the problem stated above. Owing to the complete statistical description, the information about the equilibrium states of the PS can also be obtained by reweighting methods.

However, there remains a technical problem such that a scaled long-time average, not a simple long-time average, is required to obtain a thermodynamic quantity that is defined as a space average over the phase space. In other words, the time development of the cNH cannot be interpreted as a real physical process under nonequilibrium environment with fluctuating temperature, but it may be related to an artificial development with a scaled time. The relevant problem is a theoretical complexity to construct a probability space, resulting in additional conditions for functions that defines the total distribution density.

To solve these problems, in this paper, we reconstruct an EOM that needs no scaled long time average to obtain the phase-space average. The current EOM has an invariant measure that exactly corresponds to the target density for the total system (viz., PS+TS), resulting in a simple theory, so that a simple long-time average gives the phase-space average for any phase-space function. We can monitor a real, or physically sound, development of the PS in fluctuating temperature, and find a probability that the PS should obey. Thus we can have a realistic physical process determined by the NH equation under fluctuating-temperature heat bath.

Exactly speaking, the time development of the NH equation itself may not correspond to the real physical process. However, the development should be physically sound under a small perturbation of the thermostat, and many simulation results compared with experiments support this correspondence. In this sense, the current cNH produces physically-sound time development of the target PS, which reflects the aspect of the molecular “dynamics” and should be advantageous against the Monte-Carlo “iteration” that produces unphysical time development.

In section II, we review the original cNH and consider the reason why we suffer from the scaled time average. In section III, a solution to this problem is given by reformulating a new cNH, which retains the fundamental form of the original cNH and is free from the scaled time. A relationship between the current and original cNH schemes is also demonstrated. In section IV, we propose function settings required for defining the distribution of the total system. In section V, we validate the current scheme by use of numerical simulations of simple model systems and compare the current and original schemes. We also clarify the sampling property of the current cNH by investigating how the setting of functions to define the temperature distribution affect the efficiency. Section VI concludes our study.

## II. REVIEW OF THE COUPLED NOSÉ-HOOVER EQUATION

### A. Equations of motion

The cNH equation is composed from the NH equation of PS and the NH equation of the TS. The NH equation of PS is described by the variables (*x, p, ζ*), and the NH equation of TS is described by variables (*𝒬, 𝒫, η*). Here *𝒬* is the TS coordinate, *𝒫* is the corresponding momenta, and *η* is the control variable for the TS, so that *𝒬*, *𝒫*, and *η* correspond to *x, p*, and *ζ*, respectively. In general, the number of degrees of freedom of the TS is *m*, and we denote as *𝒬* ≡ (*𝒬*_1_, …, *𝒬_m_*) ∈ ℝ^*m*^ and 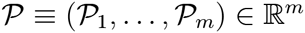, while *η* ∈ ℝ^1^ is one-dimensional as is *ζ* ∈ ℝ^1^. Thus the total phase space Ω is contained in ℝ^2(*n*+*m*+1)^ = ℝ^*n*^ × ℝ^*n*^ × ℝ × ℝ^*m*^ × ℝ^*m*^ × ℝ, and the phase-space point is denoted by 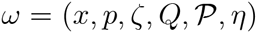. The cNH EOM, 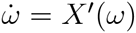, is defined as

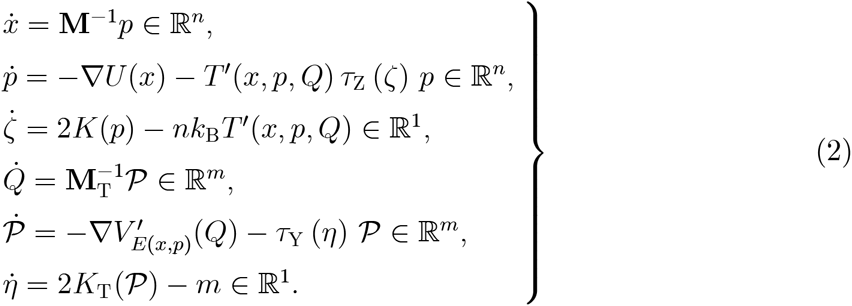

The first three equations describe the EOM of the PS and the last three describe the EOM of the TS. The PS potential energy *U* is a function on a domain *D* ⊂ ℝ^*n*^, so that Ω ≡ *D* × ℝ^*n*^ × ℝ × ℝ^*m*^ × ℝ^*m*^ × ℝ. Similarly, 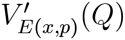 plays a role of the *TS potential energy* with respect to the coordinate *𝒬*, and it also depends on the PS energy *E*(*x,p*) ≡ *U*(*x*) + *K*(*p*). Similar to the kinetic energy of the PS, 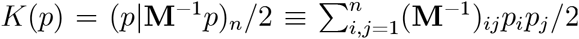, the kinetic energy of the TS is defined as

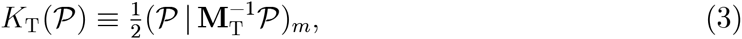

where **M**_T_ represents the *masses* for the TS. Through the last equation of (2), 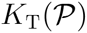 is controlled by the variable *η*, so that *η* is interpreted as a variable related to *the heat bath of the heat bath* (of the PS). Namely, the PS is under the heat bath whose temperature *T′*(*x,p,𝒬*) is dynamically changed, and the dynamics of *T′*(*x,p,𝒬*), or *𝒬*, is controlled by the TS heat bath. The functions *T′*, *τ*_Z_, *τ*_Y_, and 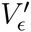 in equation (2) are defined by

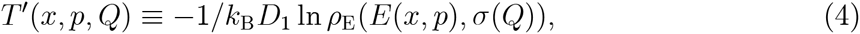

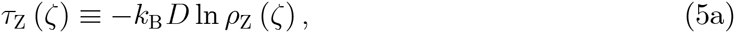

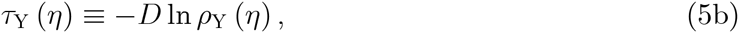

and

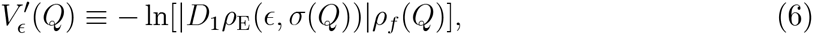

with

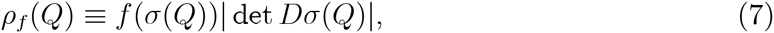

where *D* denotes the differentiation and *D_a_* denotes the partial differentiation with respect to the *a*th variable.

Physical interpretation of the functions, *σ*, *ρ*_E_, *ρ*_Z_, *ρ*_Y_, and *f*, appearing in equations (4)–(7) and their motivation of the introduction are as follows. *σ* is a map from ℝ^*m*^ into ℝ^*m*^, and we suppose that *γ* ≡ *σ*(*𝒬*) ∈ ℝ^*m*^ represents a variable that is originally a parameter and will be treated as a dynamical variable. The reason why we use the function *σ*, instead of just *γ*, is to easily handle the domain (accessible area) Γ for *γ*, since Γ is not necessarily the whole domain ℝ^*m*^ [e.g., if *γ* ≡ *β* ∈ ℝ, which is the (inverse) temperature, then *β* > 0 is required, so that 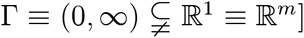] and to to easily establish an ODE on ℝ^2(*n*+*m*+1)^ [while the flow generated by an ODE is not easily handled on a restricted region of ℝ^2(*n*+*m*+1)^]. *ρ*_E_ is a function of both the PS energy *E*(*x,p*) and the parameter *γ*, viz., (*ϵ, γ*) ↦ *ρ*_E_(*ϵ, γ*). For example, *ρ*_E_(*ρ,β*) ≡ exp [−*βϵ*], by putting *m* = 1, represents the usual Boltzmann-Gibbs (BG) density, and so *γ* ≡ *β* describes the inverse temperature. We here mainly consider this case, and then equation (4) yields

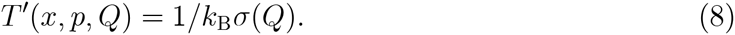

Namely, the *functional temperature T*(*x, p,𝒬*) acting on the PS really becomes the *dynamical (heat-bath) temperature* 1/*σ*(*𝒬*), which is originally a constant temperature parameter 1/*β*. *ρ*_z_(*ζ*) and *ρ*_Y_ (*η*) denote distribution functions of the control variables for PS and TS, respectively, and typically we use Gaussian forms,

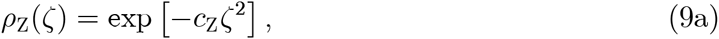

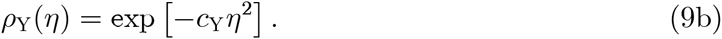

Finally, *f*: *σ*(ℝ^*m*^) → ℝ describes the distribution of *γ* ≡ *σ*(*𝒬*) and governs the statistical feature of the dynamical parameter *γ* or dynamical inverse temperature 1/*k*_B_*σ*(*𝒬*).

### B. Distribution density

These functions *σ*, *ρ*_E_, *ρ*_Z_, *ρ*_Y_, and *f* are relating to a function *ρ*, which is a underling density of the EOM, as discussed below. *ρ* is explicitly described as

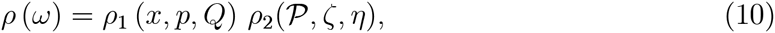

where

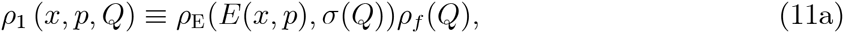

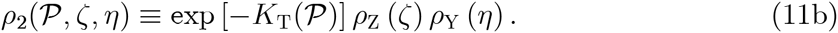

This *ρ* will be related to a smooth density of an invariant measure for the flow generated by the ODE (2). To ensure this issue in the original scheme explained in this section and to consider a new scheme presented in section III, we here summarize mathematical conditions for these functions and related quantities, as follows:

#### Condition 1

*σ*: ℝ^*m*^ → ℝ^*m*^ *is a C*^3^-*diffeomorphism, and* det *Dσ*: ℝ^*m*^ → ℝ *has a definite signature. ρ*_E_: ℝ × ℝ^*m*^ ⊃ *O* × *σ*(ℝ^*m*^) → ℝ_+_ *is of class C*^2^, *where O is an open set containing E*(*D* × ℝ^*n*^) *[ℝ_+_ denotes strictly positive real numbers]. U is a C*^2^ *function on a domain D* ⊂ ℝ^*n*^. **M** *and* **M**_*T*_ *are symmetric, positive-definite square matrices. ρ*_Z_, *ρ*_Y_: ℝ^1^ → ℝ_+_ *are of class C*^2^ *and (Lebesgue) integrable functions. f*: *σ*(ℝ^*m*^) → ℝ_+_ *is of class C*^2^ *and integrable. Assume that D* × ℝ^*n*^ × *σ*(ℝ^*m*^) → ℝ_+_, (*x,p,β*) ↦ *ρ_E_*(*E*(*x,p*), *β*)*f*(*β*), *is integrable*.

Note that additional conditions are further required for the original cNH scheme in order to validate the scaled-time average, as detailed in [12].

### C. The scaled-time average

The EOM (2) realizes the density in the sense that the following relation holds under an ergodic assumption:

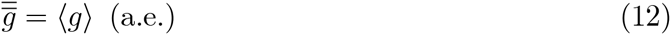

Here the right-hand side (RHS) is the space average under the density *ρ*,

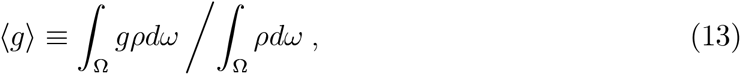

where *g*: Ω → ℝ is a phase-space map satisfying ∫_Ω_ |*gρ*|*dω* < ∞, and *dω* is the Lebesgue measure on ℝ^2(*n*+*m*+1)^. The left-hand side (LHS) of equation (12) is a “scaled-time” average defined by

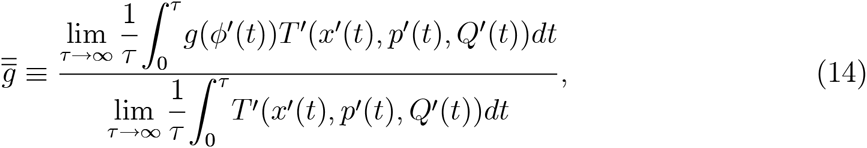

where 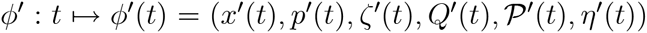 is a solution of 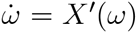. If we use, instead of the scaled time average 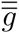, the following simple time average

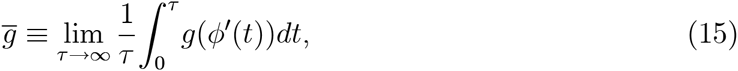

then we have

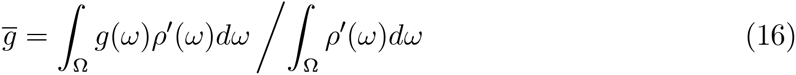

for almost all initial value of *φ′*, where *ρ’* denotes a scaled density function defined by

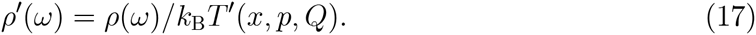

Here, equation (16) is due to the equilibrium Liouville equation

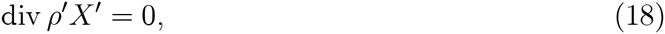

which ensures that *ρ′dω* becomes an invariant measure of the flow of *X′*. Equations (16) and (12) are obtained from the assumption that the flow is ergodic with respect to this invariant measure [12].

Namely, although equation (16) is valuable in that it connects the time average with the space average, the space average is defined by the scaled density *ρ’*, which is not the target density *ρ*. Thus we have developed the scaled-time average, so that we have obtained equation (12). However, as stated in section I, the scaled-time average is not convenient when we observe the dynamics of the ODE, because the time development should not straightforwardly correspond to the realistic physical process for producing *ρ*. The current study is motivated to solve this problem.

### D. Joint distribution for *x, p*, and *β*

According to equation (12), for a function of the form *g*(*ω*) = *B*(*x,p, σ*(*𝒬*)) = *B*(*x,p, β*), we have [12]

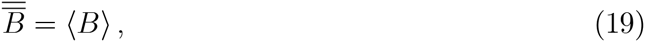

where

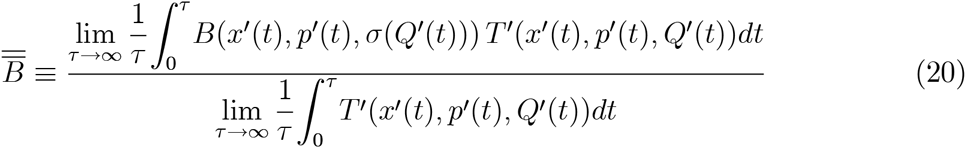

and

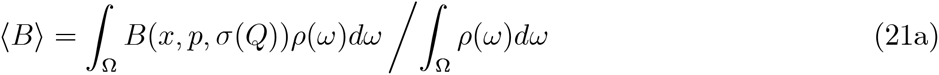

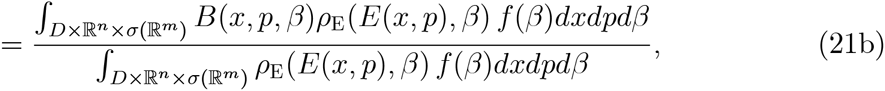

assuming that *B* gives a finite numerator of equation (21b). Namely, (*x,p, β*) is generated with the probability distribution *β*_E_(*E*(*x,p*),*β*) *f* (*β*)*dxdpdβ*, which means that *β* is fluctuating according to the distribution density function *f*. This is the critical point to capture the EOM. Note that | det *Dσ*(*𝒬*)| appeared in equation (7) is the Jacobian needed for the variable transformation from *𝒬* to *β* in the integrations [12, 13]. The important issue is that 〈*B*〉 retains the same form as equation (21), *even if* we will use the simple time average 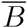, instead of 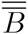, as demonstrated in the next section.

## III. COUPLED NOSÉ-HOOVER EQUATION WITH SIMPLE TIME AVERAGE

### A. General derivation

Here we obtain an EOM that is free from the time scaling so that the simple time average of the EOM yields the space average, equation (13). The aimed EOM, 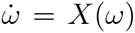, (note that we will remove the prime for quantities newly obtained) has to satisfy the Liouville equation,

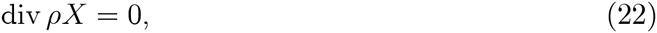

where *ρ* is defined by originally given equations (10)–(11) under Condition 1. To do this, we will take a straightforward approach, that is, to find new definitions of characteristic functions, *T′*(*x,p, 𝒬*) and 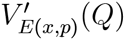, used in equation (2) such that they contribute to the Liouville equation. Namely, using new notations, we specifically consider the following EOM and seek *T*(*x,p, 𝒬*) and *V*(*x,p,𝒬*) to meet equation (22):

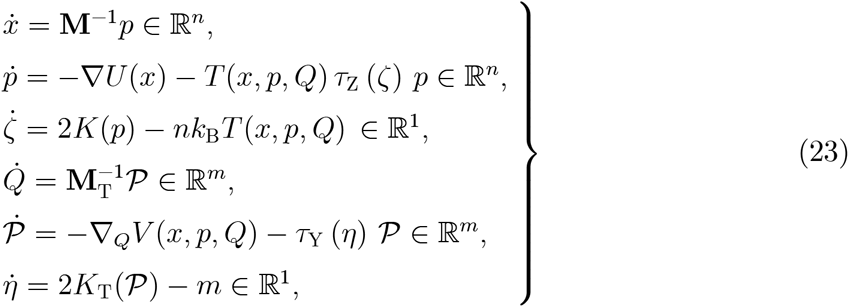

where *τ*_Z_(*ζ*), *τ*_Y_ (*η*), and 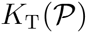 are the ones previously defined. Here *T* and *V* are assumed to be smooth (classes *C*^1^ and *C*^2^, respectively) so that *X* becomes *C*^1^. In addition, *T* should be strictly positive since it will denote a temperature. Note that our strategy is fundamentally similar to that of other methods for designing thermostat EOMs [17].

#### Lemma 2

*The Liouville equation (22) is equivalent to*

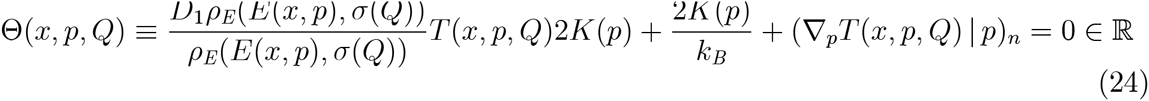

*and*

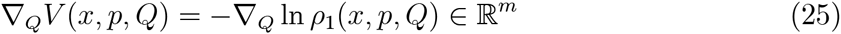

*for all* (*x,p,𝒬*) ∈ *D* × ℝ^*n*^ × ℝ^*m*^.

**Proof**. Since *ρ* > 0, equation (22) is equivalent to

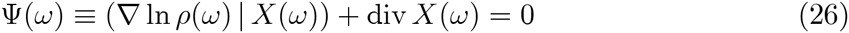

for all *ω* ∈ Ω. By a straightforward calculus we have

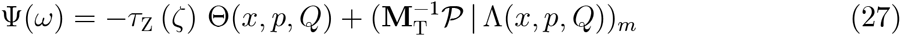

for any *ω*, where

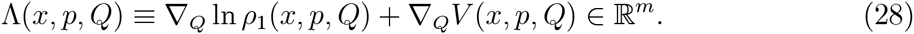

We easily see that Θ = 0 and Λ = 0 give Ψ = 0. We also show the converse, as follows: Since *τ*_Z_ defined by equation (5a) is *C*^1^ (*ρ*_Z_ is *C*^2^) and since *ρ*_Z_ is integrable, there exists *ζ*_0_ such that *Dτ*_Z_ (*ζ*_0_) ≠ 0. Thus, for any (*x,p,𝒬*) [and for any 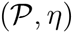], 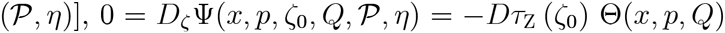, so that Θ = 0. Hence, for any (*x,p,𝒬*) and for any *l* = 1, …, *m*, taking 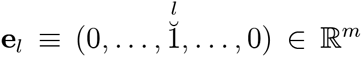, we have 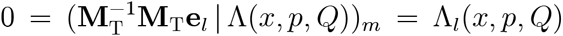. Thus Λ = 0. Therefore, the validity of equation (26) for all *ω*, viz., equation (22), is equivalent to Θ = 0 and Λ = 0.

Hence, *V* is specified by equation (25) and its smoothness is consistent with that of ln *ρ*_1_ from the assumptions. Thus the remaining task is to seek *T* that is *C*^1^, strictly positive, and satisfies equation (24). Before doing so, note that the range of *E* becomes a semi-infinite interval, i.e.,

#### Lemma 3

*E*(*D* × ℝ^*n*^) = |*α*, ∞) *for a certain α* ∈ [−∞, ∞).

**Proof** From the assumptions (*U* is *C*^0^ and *D* is connected), the range of *U* is an interval, viz., *U*(*D*) = |*ρ, β*| ⊂ ℝ, with −∞ ≤ *α* < *β* ≤ ∞. Here *∞* ≠ *β*, because *U* should not be constant; otherwise, *β* is irrelevant to *x* and thus the integrability condition fails. From the assumption of **M**, the range of *K* is a semi-infinite interval, i.e., *K*(ℝ^*n*^) = [0, ∞). Thus *E*(*D* × ℝ^*n*^) = {*u* + *k* | *u* ∈ |*α, β*|, *k* ∈ [0, ∞)} = {*α*, ∞), where the left-end type of the range corresponds to that of *U*(*D*).

In what follows, we assume *U*(*D*) = (*α*, *β*|, as encountered in many applications. Thus *E*(*D* × ℝ^*n*^) = (*α*, ∞) =: *J_E_*, which simplifies the below discussion.

To seek *T*(*x,p,𝒬*), we assume a form that is similar to *ρ*_E_, viz., 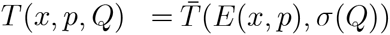 for a certain function 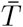. Under this assumption we can completely describe the feature of 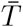, which is proved in an elementary fashion given as follows [note that *k*_B_ > 0 in equation (24) can be dropped, using *T* instead of *k*_B_*T*]:

#### Proposition 4

*Let* 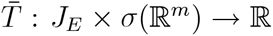 *be of class C*^1^. *Let ρ*_E_: *J_E_* × *σ*(ℝ^*m*^) → ℝ *satisfy* ∫_(*ϵ*_0_,∞)_ *ρ_E_*(*ϵ, β*)*dϵ* < ∞ *for any* (*ϵ*_0_,*β*) ∈ *J_E_* × *σ*(ℝ^*m*^). *Define T*: *D* × ℝ^*n*^ × ℝ^*m*^ → ℝ, 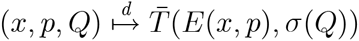, *which becomes C*^1^. *Then, the condition of T such that*

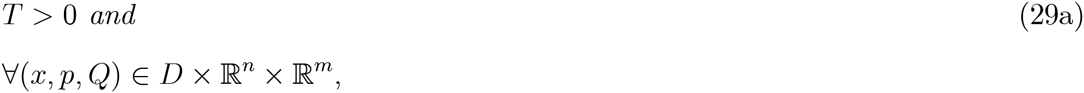

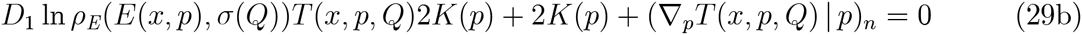

*is equivalent to the condition of* 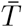 *such that*

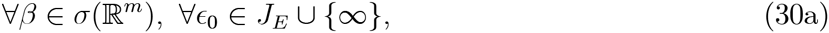

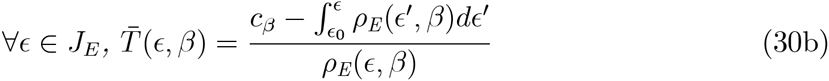

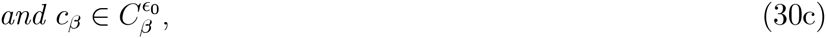

*where* 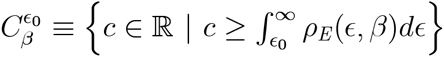.

**Proof** First, for equation (30), it follows from the assumptions of *ρ*_E_ that 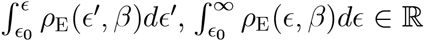 holds, so that 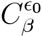 is not empty, for any *ϵ*_0_ ∈ *J_E_* ∪ {1∞} and any (*ϵ, β*) ∈ *J_E_* × *σ*(ℝ^*m*^).

Now, the positivity condition *T* > 0 in equation (29a) can be replaced with

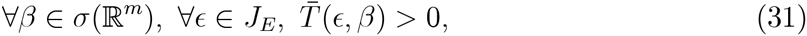

using the fact that *E*: *D* × ℝ^*n*^ → *J_E_* is onto, due to Lemma 3 (with the open interval assumption). For the other condition in equation (29), first we see that *T* becomes *C*^1^ and 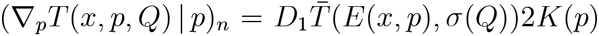 for all (*x,p, 𝒬*) ∈ *D* × ℝ^*n*^ × ℝ^*m*^. Thus equation (29b) is equivalent to

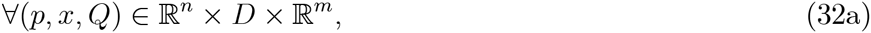

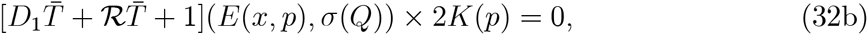

where 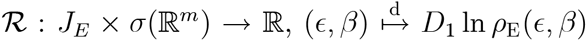. Note that *K*(*p*) = 0 if only if *p* = 0. Since the map from ℝ^*n*^ × *D* × ℝ^*m*^ to ℝ defined by the LHS of equation (32b) is continuous, we can ignore the point *p* = 0. Namely equation (32) is equivalent to

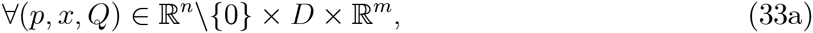

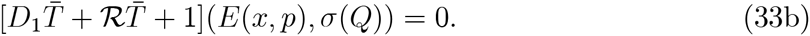

Again, from the continuity of the map defined by the LHS of equation (33b), we can add the point *p* = 0 in equation (33). Hence, equation (29b) is equivalent to

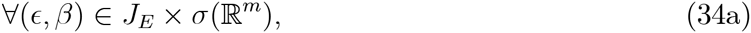

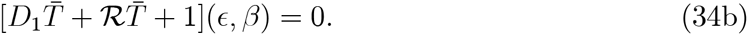

While partial derivative is used, this yields essentially an ODE with respect to 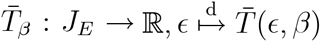 for each *β* ∈ *σ*(ℝ^*m*^). Namely, equation (34) is equivalent to

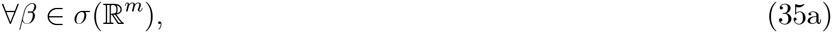

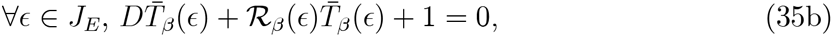

where 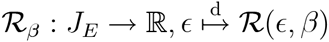.

Below, we fix an arbitrary *β* ∈ *σ*(ℝ^*m*^) and concentrate on equation (35b). Since equation (35b) is a one-dimensional linear ODE having a continuous coefficient 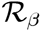, its solution satisfying an "initial" condition 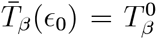 for any *ϵ*_0_ ∈ *J_E_* and any 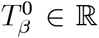 is uniquely given as

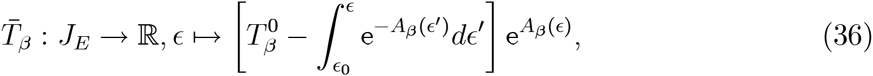

with

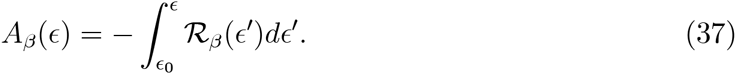

Going back to the definition of 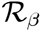 and defining 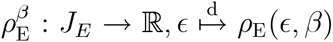 (for any *β*), we have

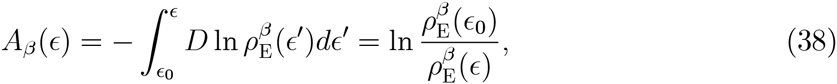

so that 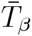 defined by equation (36) becomes

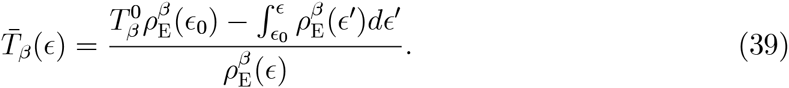

Hence, we can show that equation (35b) and a positivity condition that comes from equation (31), viz.,

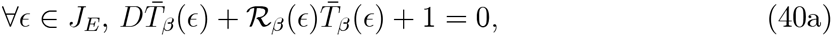

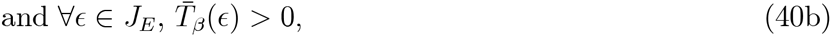

are equivalent to

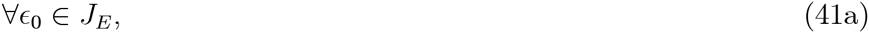

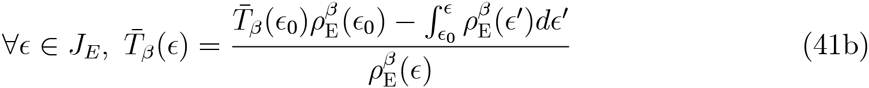

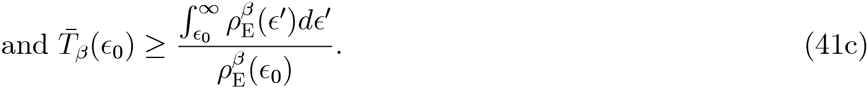

We first show equation (40) ⇒ equation (41). Clearly from equation (39), we see that equation (41b) is deduced for ∀_*ϵ*_0__ ∈ *J_E_*. From this and equation (40b), we have

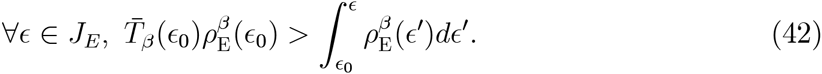

Thus 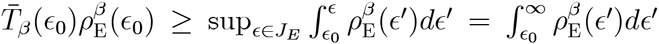, implying equation (41c). Conversely, assume equation (41). First note that equation (40a) clearly holds due to equation (41b). Second, note that equation (41c) yields 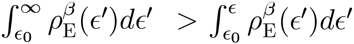 for any *ϵ* ∈ *J_E_*, indicating that the numerator of 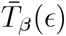 given in equation (41b) is strictly positive for ∀_*ϵ*_ ∈ *J_E_*. Thus equation (40b) is obtained.

Integrating these results, we observe that equation (29) is equivalent to the statement that

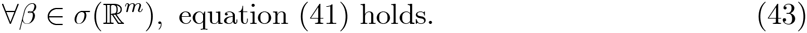

Finally we show the equivalence between equation (43) and equation (30). We can easily show equation (43) from equation (30), since it follows from a substitution of *ϵ* = *ϵ*_0_ ∈ *J_E_* into equation (30b) that 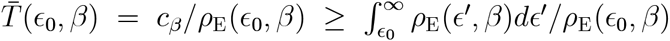. We show equation (30) from equation (43). First, consider a case in which *ϵ*_0_ ∈ *J_E_*. Define 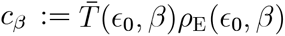, then equation (41c) implies 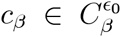, viz., equation (30c), and equation (41b) means equation (30b). Second, let *ϵ*_0_ = ∞. For an arbitrary chosen 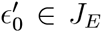, consider a substitution of 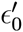 into *ϵ*_0_ of equation (41), so that equation (41b) indicates

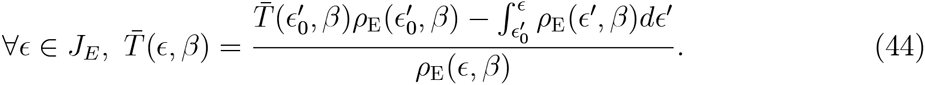

In addition, define 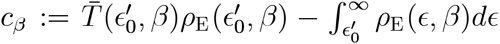. Then equation (41c) indicates *c_β_* ≥ 0, which implies 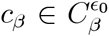 since 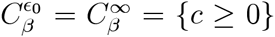. The numerator of the RHS of equation (44) becomes 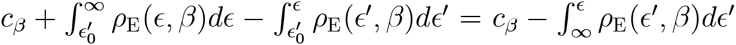, resulting in equation (30b) for *ϵ*_0_ = ∞.

Thus, the aimed EOM, 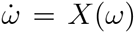, that is given by the form of equation (23) and satisfies the Liouville equation (22) is determined. The force function, −∇*_Q_V*(*x, p, 𝒬*), of the TS is completely characterized by equation (25). For the temperature function *T*(*x,p, 𝒬*), supposed to be 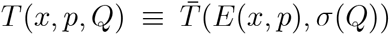, is determined, up to two constants *ϵ*_0_ ∈ *J_E_* ∪ {∞} and 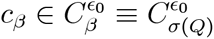, by

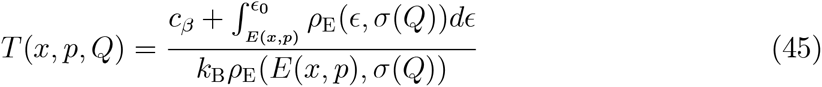

for all (*x, p, 𝒬*) ∈ *D* × ℝ^*n*^ × ℝ^*m*^. Regarding the constant *c_β_* (which has an energy dimension), its range 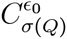 depends on *𝒬* in general, which may be inconvenient in applications. An exceptional case is of *ϵ*_0_ = ∞, for which 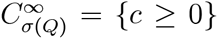 for any *𝒬*, indicating that any positive constant serves as *c_β_*. Thus we will use this choice.

### B. Resulted forms

Therefore, the new *coupled NH* EOM, 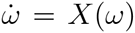, is represented by equation (23) with the following two function forms: (I) The TS potential represented as

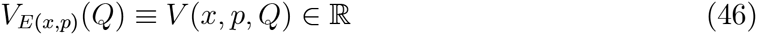

is defined by

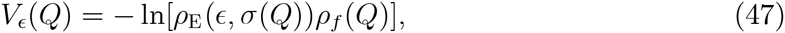

which yields the TS force − ∇*V*_*E*(*x, p*)_(*𝒬*) = − ∇*_Q_V*(*x,p,𝒬*) ∈ ℝ^*m*^; (II) the dynamical temperature is defined as

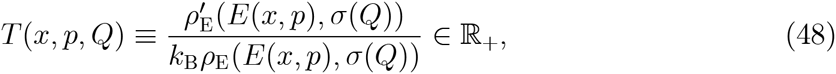

where

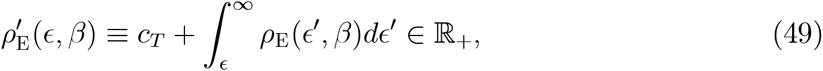

with any constant *C_T_* > 0.

The integration of *ρ*_E_ is needed in the current scheme, but it should not be the bottleneck. This is because the integral is one dimensional, so that an explicit integration can be done for a simple function or a numerical integration can be accurately performed in general.

*Remark*. The current scheme is applicable to the density form defined by equation (11a). A more general case, i.e., a use of *ρ*_Phys_(*x,p,𝒬*) considered in [12], instead of *ρ*_E_(*E*(*x,p*), *σ*(*𝒬*)), is beyond the current target.

### C. Simple time average

From the Liouville equation and the assumption that *X* is complete, its flow is invariant with respect to the measure *P* = *ρdω*, where *ρ* is given by equations (10)–(11). Thus, the probability space is simply constructed from 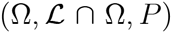, where 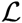 represents the Lebesgue measurable sets on ℝ^2(*n*+*m*+1)^ [18, 19]. Hence, if the flow is ergodic with respect to the measure space 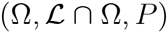, then the long-time average of any (Borel measurable) *P*-integrable phase-space function *g*, for any solution *ϕ* of equation (23), gives (for a.e. *ω*) the space average under the density *ρ*:

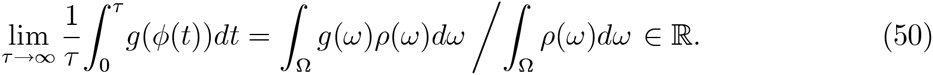

Namely, *ρ* can be realized without the scaled-time average, but with the simple time average. Thus our aim has been attained, and the cNH EOM uses a “real” (physically realistic) time [4].

In particular, equation (19) is replaced by

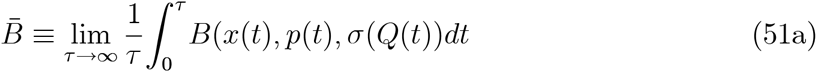

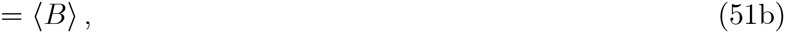

where 〈*B*〉 is given by equation (21). This indicates that the distribution *ρ*_E_(*E*(*x,p*), *β*) *f*(*β*)*dxdpdβ* is realized with the simple time average. In addition, for a physical quantity *A*: *D* × ℝ^*n*^ → ℝ, by taking a similar procedure discussed in [12, 13], we have

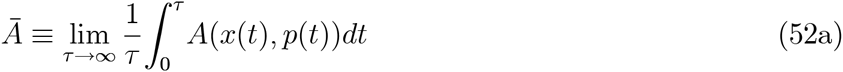

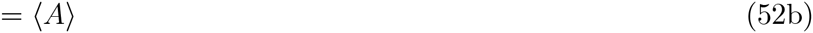

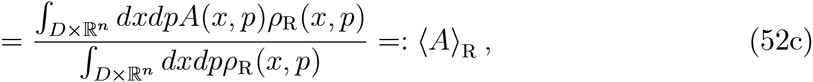

where

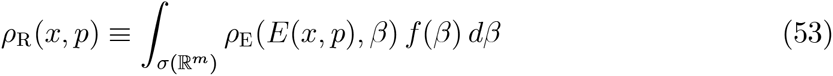

is the (unnormalized) marginal distribution of (*x,p*). We have assumed that the numerator of equation (52c) is finite.

A reweighting to any other density *ρ*_TRG_: *D* × ℝ^*n*^ → ℝ_+_ is possible by a reweighting formula

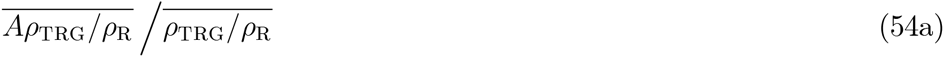

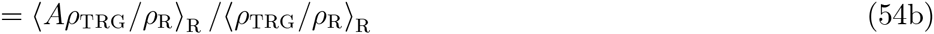

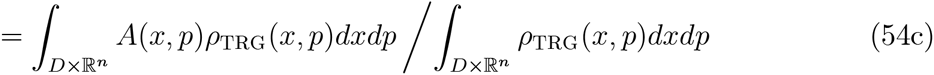

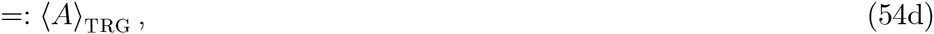

provided that the space averages in equation (54c) are finite. An alternative reweighting formula that needs no information on *ρ*_R_ can be obtained [12] as

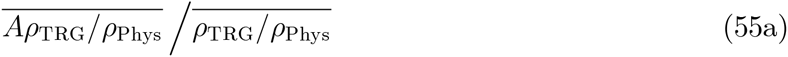

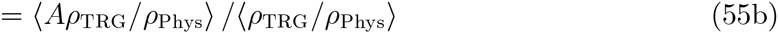

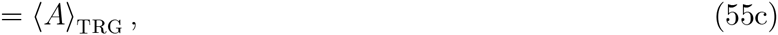

where 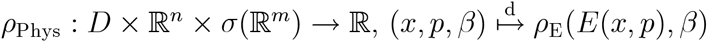.

### D. Relationship with the original scheme

The current scheme is intimately related to the original scheme. To see this, consider 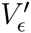 and *T′* that are originally given by equations (6) and (4), respectively, and the current ones *V_ϵ_* and *T* that are given by equations (47) and (48), respectively. Then, notice that 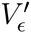 and *T′* with using 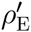 [equation (49)], instead of *ρ*_E_, are equivalent to the current *V_ϵ_* and *T*, respectively. This can be seen, using a relation

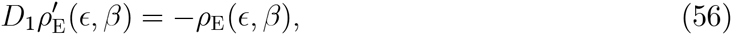

as

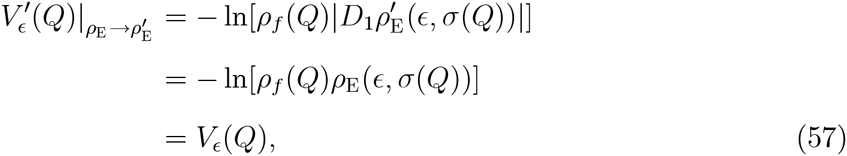

and

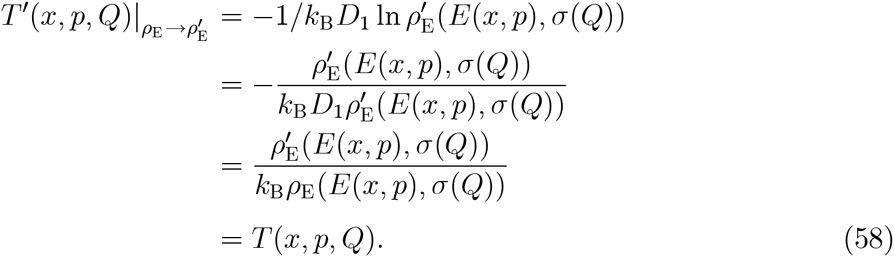

Thus the current scheme is equivalent to *the original scheme with using* 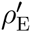, instead of *ρ*_E_. Exactly speaking, before this claim is valid, we should state that the signature conditions for density 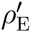 required in the original scheme are met, i.e., 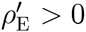 and 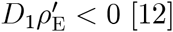 [12]. These can be ensured by equations (49) and (56), respectively.

Second, we see that the scaled density,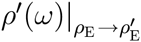, which is equation (17) using 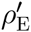 instead of *ρ*_E_, becomes

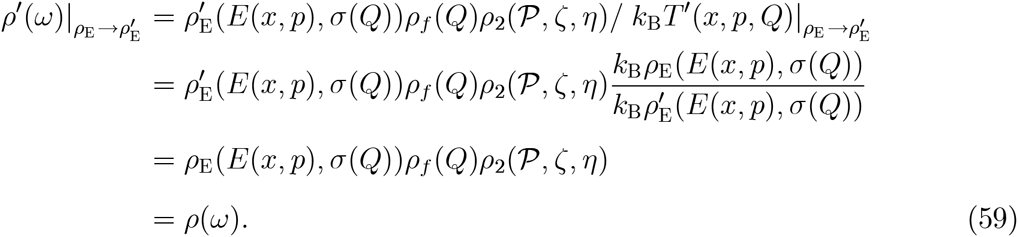

Namely, 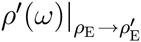 is equivalent to the target *ρ*(*ω*).

This fact leads to an alternative derivation of the aimed equation (50). That is, if we use the original scheme with 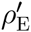, *instead of ρ*_E_, then the combination of the original-scheme formula equation (16) and the density-relationship equation (59) give

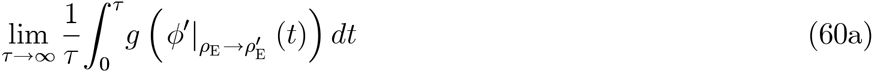

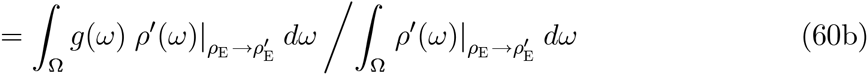

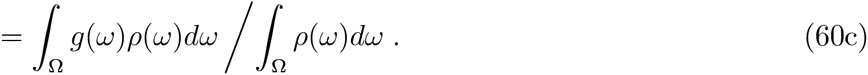

Here, we see that 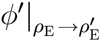 in the LHS is nothing but the solution of the current scheme, due to equations (57) and (58). Thus it can be denoted as *φ*, and equation (50) is derived.

## IV. FUNCTION SETTING

In applications we should set functions, *ρ*_E_, *σ*, and *f*. First we consider *ρ*_E_, and then we give two examples for *σ* and *f*. The first example, given in section IV B, was also used for the original scheme, and we reconsider a relationship between the original and current schemes with this function setting. The second example for σ and f is given in section IV C, as a new example.

### A. Boltzmann-Gibbs *ρ*_E_

As a physically important example, we define *ρ*_E_ by the Boltzmann-Gibbs distribution, using *m* = 1, as

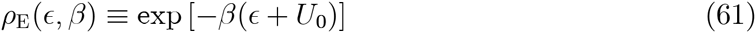

for (*ϵ, β*) ∈ *J_E_* × *σ*(ℝ^1^), with *σ*(ℝ^1^) ⊂ ℝ_+_ being assumed. Here *U*_0_ is an adjustable parameter [12, 13].

Then, the conditions stated in Condition 1 and Proposition 4 are met, and the current scheme gives

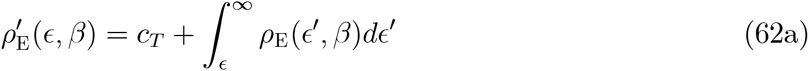

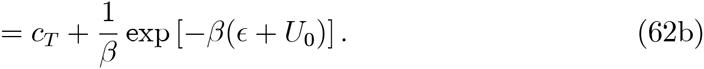

In this case we get

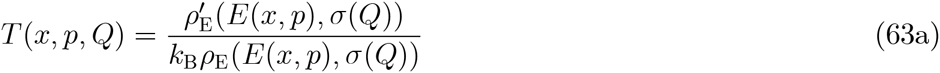

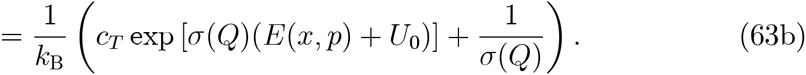

### B. Gamma *f* with exponential *σ*

As presented in the original scheme, we can define exponential *σ* as

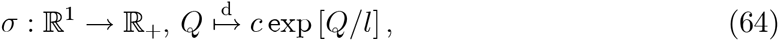

where *c, l* > 0 are parameters, and define the (normalized) gamma distribution (having parameters *α*_1_, *α*_2_ > 0)

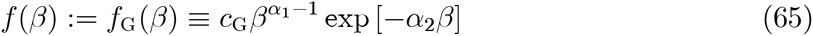

for any *β* ∈ *σ*(ℝ^1^) = ℝ_+_.

Combination of these functions with equation (61) gives

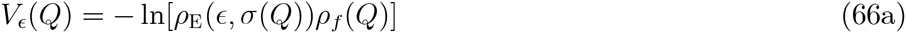

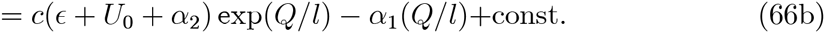

in the current scheme. Note that *f*_G_(*β*) produces the the Tsallis distribution [20–22] with the renormalized temperature *T*_0_ ≡ *α*_2_/*α*_1_*k*_B_ and the nonextensive parameter *q* ≡ 1 + 1/*α*_1_: ∫_(0,∞)_ e^−*β*(*E*(*x,p*)+*U*_0_)^ *f*_G_(*β*) *dβ* = [1 − (1 − *q*)(*E*(*x,p*) + *U*_0_)/*k*_B_*T*_0_]^1/(1−*q*)^. Note also that *V_ϵ_*(*𝒬*) becomes the Toda potential [23] for *𝒬*, representing an anharmonic spring interaction.

#### 1. Resulting relationship between the two schemes

Relationship between the original and current schemes under a use of the three functions, defined by equations (61), (64), and (65), is considered. In this case, we see that the difference between the two schemes is small to be just parameter differences. In fact, the original scheme with these three equations gives [12, 13]

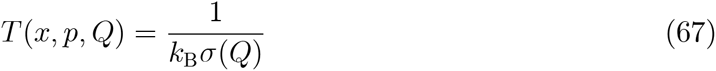

and

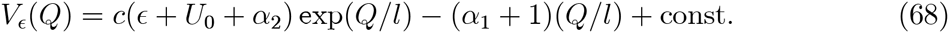

The difference between equations (63) and (67) comes from the existence of *c_T_*, and the difference vanishes if *c_T_* = 0. The difference between equations (66) and (68) is just *α*_1_ (except the additive constants irrelevant to the EOM), and the latter is obtained if *α*_1_ + 1 is used, instead of *α*_1_, in the former. In other words, with these function definitions, the EOM (dynamics) of the original scheme with *α*_1_ − 1 is equivalent to EOM (dynamics) of the current scheme with *α*_1_ and *c_T_* = 0. Thus the trajectories of any variables obtained by the original scheme (note that all applications in the original scheme employed these three functions) can be interpreted to be physically realistic via the redefinition of the parameter value, since the those of the current scheme are physically realistic.

However, the distribution obtained by the original scheme with *α*_1_ − 1is not equivalent to the distribution obtained by the current scheme with *α*_1_ and *c_T_* = 0 in general (the marginal distributions of *x, p*, and *β* are different due to the difference in *f*, but the marginal distributions of 𝒫, *ζ*, and *η*, which are irrelevant to *f*, are the same eventually). In contrast, between the original scheme with *α*_1_ and the current scheme with *α*_1_ and *c_T_* = 0, the EOMs (dynamics) are different, but the distributions are the same (although the distribution for the original scheme is generated by the scaled time average and the distribution for the current scheme is generated by the simple time average). These issues are also numerically confirmed in the next section.

### C. Beta *f* with sigmoid *σ*

*σ* governs the range of the dynamical parameter *σ*(*𝒬*), as stated in section II. Since *σ*(*𝒬*) employed in *ρ*_E_ defined by equation (61) represents the inverse temperature, it is possible to freely set the infimum and supremum of the inverse temperature in the current method [note that the dynamical temperature, which fluctuates and acts on the PS, becomes *T*(*x,p, 𝒬*) = 1/*σ*(*𝒬*) due to equation (63) if *c_T_* = 0]. For this purpose, under equation (61), we can use a sigmoid function for *σ*:

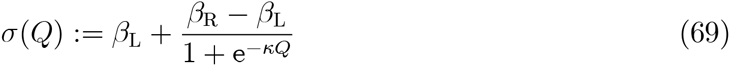

with parameters *κ* > 0 and *β*_R_ > *β*_L_ ≥ 0. This function meets the requirements in Condition 1 and yields *σ*(ℝ^1^) =]*β*_L_, *β*_R_[=: *I*_B_, viz., *β*_L_ is the infimum and *β*_R_ is the supremum. Note that a limiting case of *β*_L_ → 0 might be viewed as an extension of *σ* given by equation (64), since *σ*(*𝒬*) ~ *β*_R_*e^κQ^* as *𝒬* → −∞.

Since *f* is defined on an open interval *σ*(ℝ^1^) and since it is natural to impose decaying to 0 as *β* tends to the both ends of *σ*(ℝ^1^), the setting of *f* by equation (65) does not serve to couple with the sigmoid *σ*. For this purpose, by using the Beta distribution,

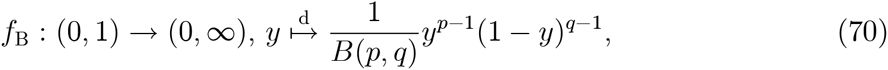

with *p, q* > 1 and 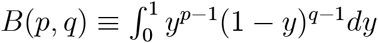, we can define *f* by

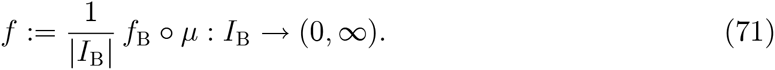

Here, 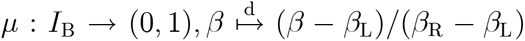 is an affine transformation so that *f* has a support on *I*_B_, and 1/|*I*_B_| ≡ 1/(*β*_R_ − *β*_L_) is the normalization constant.

In Appendix A, we detail the physical marginal distribution function, the TS potential energy, and the relationship between the NH equation and the current cNH equation yielded by equations (69) and (71).

## V. NUMERICAL EXAMINATION

To validate the current scheme, we applied it to two model potential systems, onedimensional harmonic oscillator (1HO) potential and two-dimensional Müller-Brown (2MB) potential systems. The specific purpose is to (i) see the generation of the correct distributions, (ii) numerically confirm the comparison between the current and original schemes, as done in sections IIID and IV B 1, and (iii) investigate the sampling property of the current method. Here, (i) is done in the both systems, (ii) is done in the 1HO system, which is suitable to the purpose, and (iii) is done in the 2MB system, which provides a model of a complex potential energy surface.

For numerical integration of the ODE (23), a symmetric, second-order integrator was used, which is based on the extended space formalism [24, 25] detailed in Appendix B. We integrated 10^8^ (1HO) and 10^9^ (2MB) time steps with a unit time of 1 × 10^−3^. The parameters in the EOM were as follows: PS masses **M** = **1**, TS mass **M**_T_ = 1 (*m* = 1), *c*_Z_ = *c*_Y_ = 1 [see equation (9)], *c*_T_ = 0, and *k*_B_ = 1. All quantities are treated as dimensionless.

### A. 1D harmonic oscillator system

The 1HO potential system is defined by a potential energy

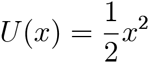

for *x* ∈ ℝ^1^ (viz., *n* = 1). Note that 1HO is a simplest but typical model that describes PS behavior around an equilibrium state. In addition, the BG distribution of the 1HO system is not easily generated by the conventional NH equation due to the lack of the ergodicity [26]. It is also convenient to compare the current and original schemes, since the latter was tested in the previous study [12].

We used the functions *ρ*_E_, *σ*, and *f* defined in equations (61), (64), and (65), respectively. The associate parameter values were the same as those used in the previous study [12], i.e., *U*_0_ = 0, *c* =1, *l* = 2.24 [to set 1 ~ (*α*_1_ + 1)/*l*^2^; see also [13]], and *α*_1_ = *α*_2_ = 4. The initial values were 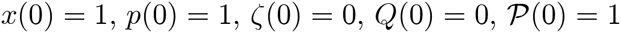, and *η*(0) = 0.

Figure 1 shows the marginal distribution densities of coordinate *x*, momentum *p*, and control variable *ζ* for the PS, and the inverse dynamical temperature *β* = *σ*(*𝒬*), momentum 𝒫, and control variable *η* for the TS. The simulated and exact (theoretical) results agree well. These results indicate that the current scheme produced a sufficiently accurate distribution. Note that although we used equations (64) and (65) for *σ* and *f* to perform the comparison below, it was found that accurate distribution densities were also obtained by using the functions of equations (69) and (71).

**Figure 1.**
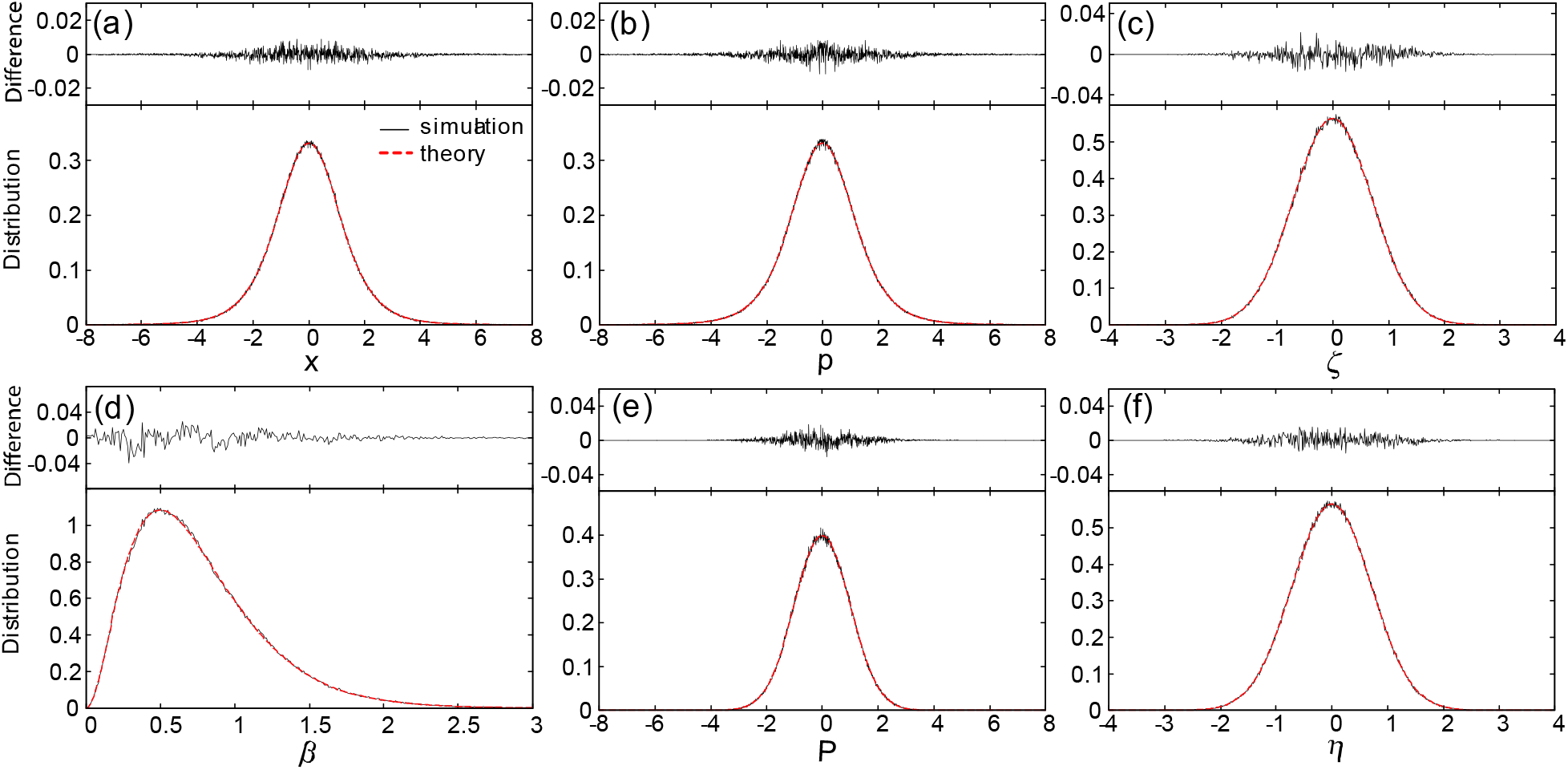
Marginal distribution densities of (a) *x*, (b) *p*, and (c) *ζ* for the physical system, and (d) the inverse dynamical temperature *β* = *σ*(*𝒬*), (e) *𝒫*, and (f) *η* for the temperature system are shown for the 1HO model. Simulated and theoretical densities are shown by solid black and dashed red, respectively, while their differences are also shown on the top.

To compare the distributions and dynamics for the original and current schemes, three simulations were carried out: (i) the original scheme with *α*_1_ =4 (the parameter of *f*), (ii) the current scheme with *α*_1_ = 4, and (iii) the current scheme with *α*_1_ = 5. As is described in section IV B1, it is expected that (i) and (ii) yield the same distribution, but different dynamics, and that (i) and (iii) yield the different distribution, but the same dynamics.

Figure 2 (a) shows the trajectory of PS coordinate *x*, physical temperature, *T*_P_ ≡ 2*K*(*p*)/*n* = *p*^2^, and the dynamical temperature, *T*_D_ ≡ *T*(*x,p, 𝒬*) = *σ*(*𝒬*)^−1^ [see equation (67)] for cases (i) and (ii) (the trajectory for case (iii) is omitted since it is the same as case (i)). We found that the dynamics are really different with each other. Nevertheless, common features of the two cases are: the trajectory of 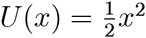 approximately keeps the original Newtonian frequency of *π* but is perturbed by the fluctuations of the temperature *T*_P_; *T*_P_ vibrates in a complicated manner but correlates with the dynamical temperature *T*_D_ derived by the TS; and thus the correlations between *T*_P_, *T*_D_, and *U*(*x*) can be seen [12, 13]. The conservation of the invariant function in the current scheme (see Appendix B) for case (ii) was good, indicating the success in the numerical integration.

**Figure 2.**
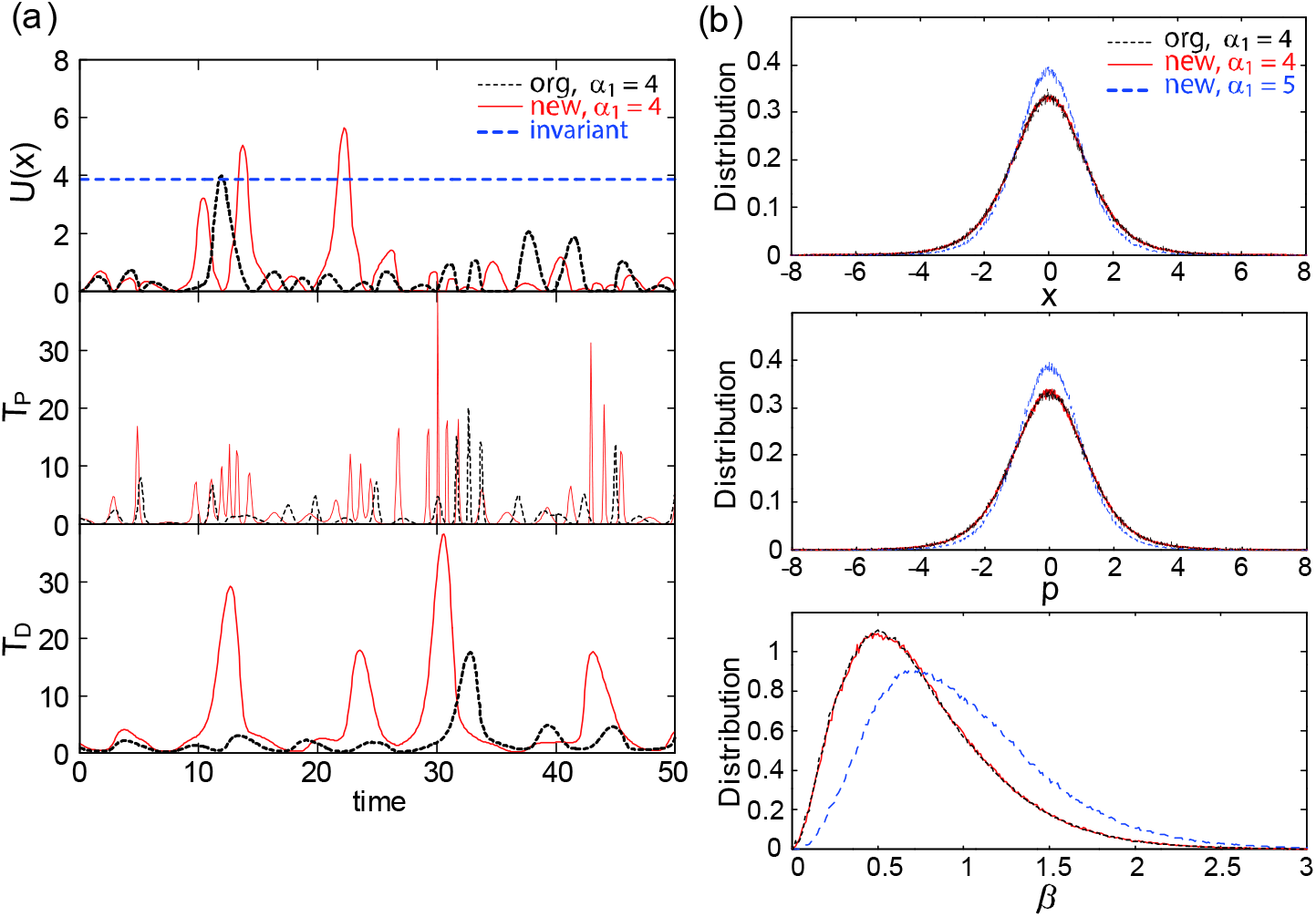
(a) Time courses of potential energy *U*(*x*), physical temperature *T*_P_ = *p*^2^ and dynamical temperature *T*_D_ = *σ*(*𝒬*)^−1^ for the 1HO model. Those simulated by the original and current cNH are shown by dotted black (case (i)) and solid red (case (ii)), respectively. Time course of invariant *L* simulated by the current cNH is also shown by dashed blue. (b) Marginal distribution densities of *x, p* and *β* = *σ*(*𝒬*) simulated by the original cNH with *α*_1_ = 4 (case (i); dotted black), and by the current cNH with *α*_1_ =4 (case (ii); solid red) and *α*_1_ =5 (case (iii); dashed blue).

Figure 2 (b) shows the distributions of the cases (i)–(iii) using *t* = 10^4^. It was clarified that (i) and (ii) generated the same distributions within a tolerance, although the dynamics are quite different, as shown in Figure 2 (a), and although the procedure to make the distribution is different, where case (i) used the scaled time average and case (ii) used the simple time average. This is really because the both schemes are accurate. We also confirmed that cases (i) and (iii) generated different distributions. The distribution of *β* for (iii) shifts right (*β* being larger) to that of (i), because the average of *β* for the 1HO system, represented by

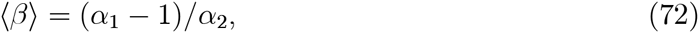

is increasing with respect to *α*_1_. These results indicate the validity of our theoretical analysis in section IV B 1.

Figure 2 (a) also shows that the current scheme (case (ii)) has larger amplitudes of *T*_D_ than the original scheme (case (i)). It seems to contradict to the fact that the two schemes with the same parameter *α*_1_ produce the same distribution. The reason really comes from the difference in making process of the distributions (scaled time average for case (i) versus the simple time average for case (ii)). Thus the trajectories, including the amplitude behavior, should be different. In fact, the dynamics of case (i) is equivalent to that of case (iii), and equations (72) and (51) indicate 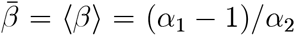, so that case (iii) (current scheme with *α*_1_ = 5) takes larger values of *β*, viz., smaller values of *T*_D_ = *σ*(*𝒬*)^−1^ = *β*^−1^, compared with case (ii) (current scheme with *α*_1_ =4). This explains the result that the *T*_D_ amplitudes in case (i) are smaller than that in case (ii).

### B. 2D Müller-Brown potential system

The Müller-Brown potential [27] is defined by

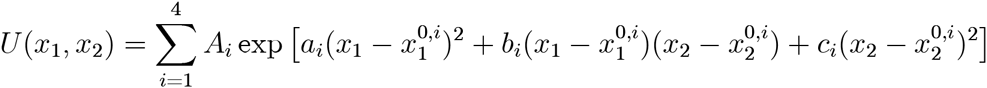

for (*x*_1_, *x*_2_) ∈ ℝ^2^ (viz., *n* = 2). The parameters used here were *A*_1_ = −20, *A*_2_ = −10, *A*_3_ = −17, *A*_4_ = 1.5, and the values of the other parameters, 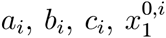, and 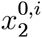, were the same as those in [27]. *U*(*x*_1_, *x*_2_) has three minima contributed by *i* = 1, 2, 3 (so *A*_1_, *A*_2_, and *A*_3_ are relevant to potential-well depth, and the other parameters are relevant to the well shapes and locations) along with two saddle points, and grows as ║*x*║ → ∞ via the contribution of *i* = 4; see figure 3(a). This potential function has been used as a model system for the studies of conformational sampling, minimum free-energy path, and path sampling [28–30]. The initial values for the cNH equation were 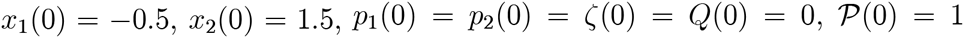, and *η*(0) = 0. Note that the initial PS coordinate was located close to the deepest minimum of *U*.

**Figure 3.**
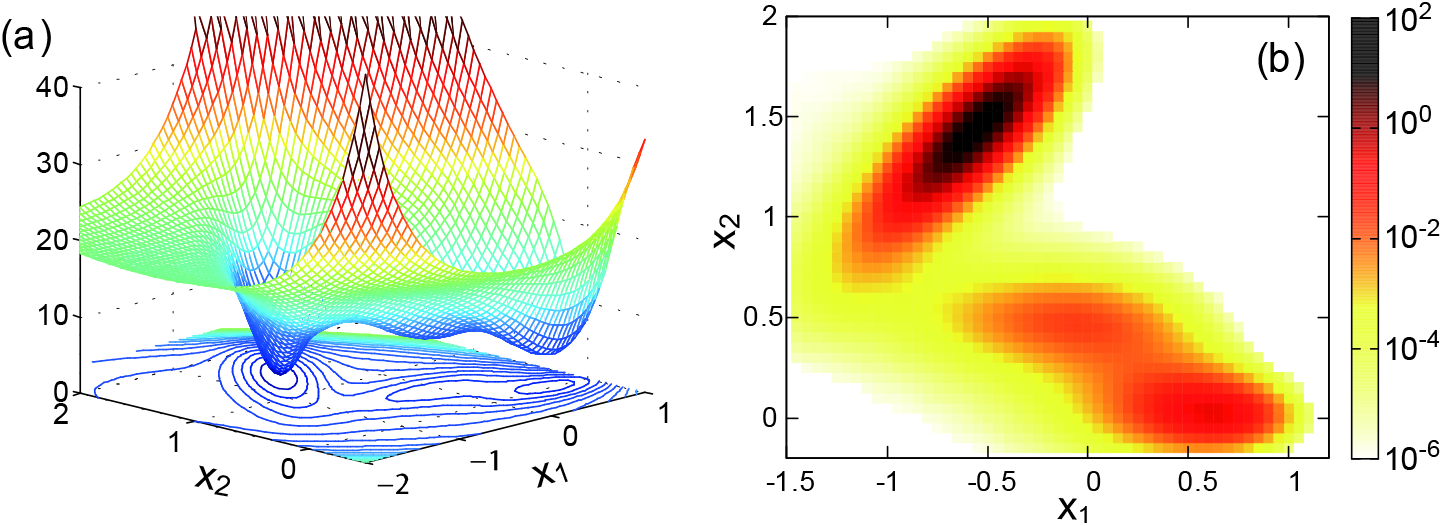
(a) The surface and contour for the 2D Müller-Brown potential *U*(*x*_1_, *x*_2_). (b) 2D distribution density with respect to *x*_1_ and *x*_2_ after reweighting to the BG distribution at temperature *T*_BG_ = 1.

We used equation (61) for *ρ*_E_ with *U*_0_ = 14.67 (which realizes an optional setting, *U*+*U*_0_ ≳ 0) and used equations (69) and (71) for *σ* and *f*, respectively. This was to investigate the effect of temperature range on the sampling performance of the current method. Although *σ* is not used in the final expression of the space average, as seen in equations (21) and (51), *σ* appears in the EOM (23) so that it should affect the sampling ability. In the present simulations we put *κ* = 1 and *p* = *q* = 5, and varied the range of ft by changing *β*_L_ and *β*_R_.

We calculated the probability density, *ρ*_BG_(*x*), for coordinates *x* = (*x*_1_, *x*_2_) under the BG distribution *ρ*_BG_ (*x, p*)*dxdp*/*Z*_BG_ ≡ e^−*β*_BG_*E*(*x,p*)^*dxdp*/*Z*_BG_ with *β*_BG_ = 1. Using the trajectory generated by the current method with *β*_L_ = 0.05 and *β*_R_ = 2, the simulated value was derived by the reweighting formula, equation (54), by the substitution of *ρ*_TRG_ ≡ *ρ*_BG_ and *A*(*x,p*) = *χ_I_*(*x*) (where *χ_I_*(*x*) = 1 if *x* ∈ *I* and *χ_I_*(*x*) = 0 otherwise) with *I* being each small bin. Figure 3(b) shows the density *ρ*_BG_(*x*), indicating full coverage of the three minima and the connecting paths. The simulated and the exact (theoretical) *ρ*_BG_(*x*) agreed well, as was seen in the distribution error of 9.2 × 10^−3^, indicating a sufficiently accurate sampling of the PS coordinates.

Figure 4(a) depicts the simulated distributions of *β* when using (*β*_L_, *β*_R_) = (0.5, 2) and (*β*_L_, *β*_R_) = (0, 5), as well as (*β*_L_, *β*_R_) = (0.05, 2), indicating the realization of the distributions as suggested by the inputs of *f*(*β*). To examine the relation between the sampling efficiency and the *β* distribution, governed by *β*_L_ and *β*_R_ (which regulate the range of *β*), we calculated the errors for the BG distribution with *β*_BG_ = 1 obtained at time *t* for both *x*_1_ and *x*_2_, Δ_*x*_1__(*t*) and Δ_*x*_2__(*t*). Here, 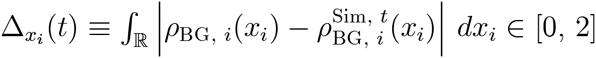, with *ρ*_BG_, *i*(*x_i_*) being the theoretical marginal distribution density for *ρ*_BG_(*x*_1_, *x*_2_) with respect to *x_i_*, and 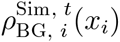 being the corresponding simulated density obtained at time *t*. Figures 4(b) and 4(c) show that the convergence is the fastest for (*β*_L_, *β*_R_) = (0.05, 2), where Pl is sufficiently small or *T*_D_ = *σ*(*𝒬*)^−1^ = *β*^−1^ becomes large so as to cross the energy barriers among the three minima, and where the *β* distribution has a relatively large density at the target inverse temperature *β* = 1 = *β*_BG_. In contrast, the convergence for (*β*_L_, *β*_R_) = (0.5, 2) is slower because *β*_L_ is too large or *T*_D_ is too small to cover the whole 2D range quickly. That for (*β*_L_, *β*_R_) = (0, 5) is slightly slower due to the smaller density of the *β* distribution at the target inverse temperature. In sum, efficient sampling can be established by using small *β*_L_ as to cover the important region relevant to jumps among stable minima, and by using not so large *β*_R_ so as to decrease the range of *β* distribution and maximize the *β* distribution density around the target inverse temperature.

**Figure 4.**
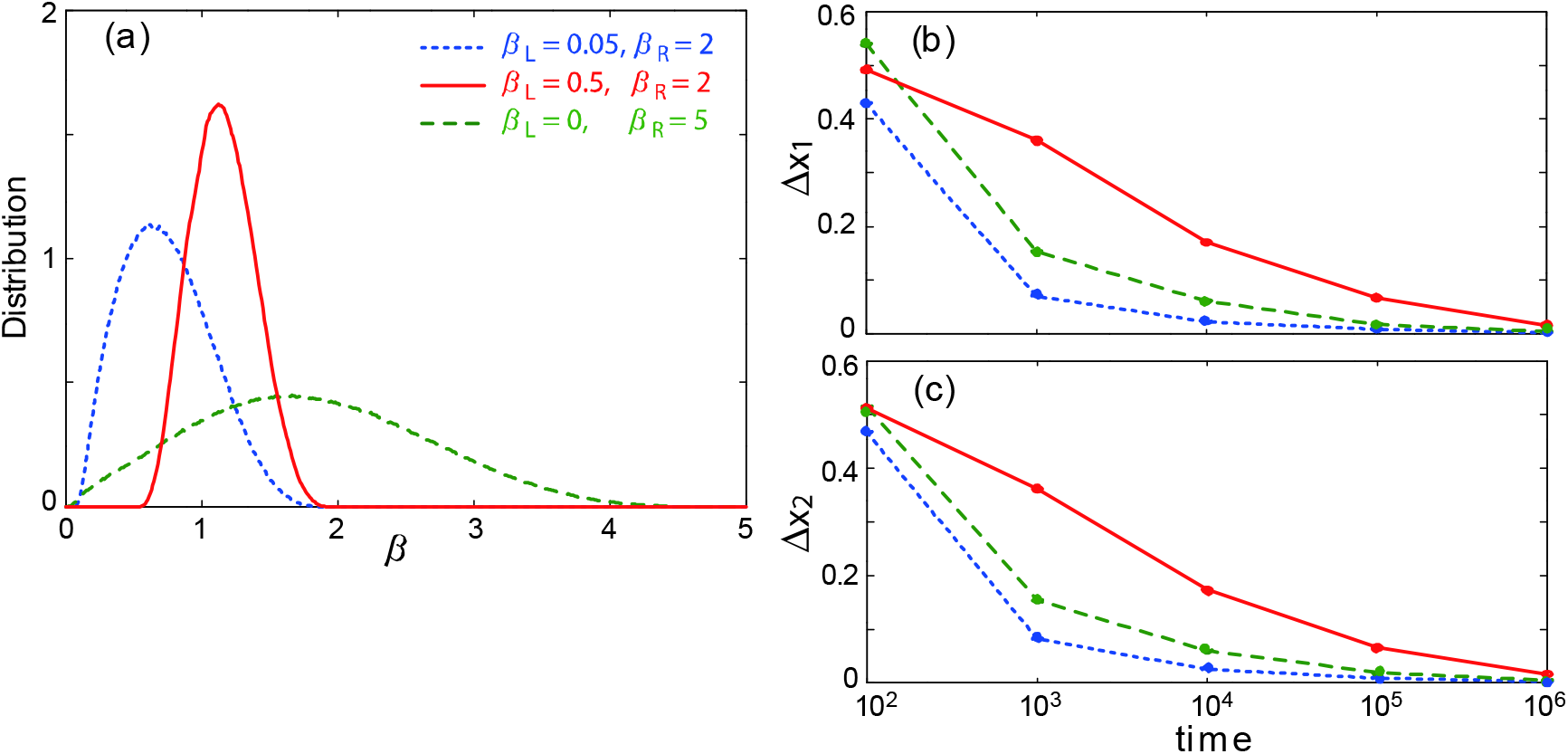
(a) Marginal distribution densities of *β* = *σ*(*𝒬*) for the 2D model. Differences between simulation and theory in the 1D marginal distributions for (b) *x*_1_ and (c) *x*_2_ after reweighting to the BG distribution at *T*_BG_ = 1, Δ_*x*_1__(*t*) and Δ_*x*_2__(*t*), respectively, are shown as a function of the simulation time *t*. The parameters in the cNH is *β*_L_ = 0.05/*β*_R_ = 2 (dotted blue), *β*_L_ = 0.5/*β*_R_ = 2 (solid red), and *β*_L_ = 0/*β*_R_ =5 (dashed green).

Figure 5(a) shows the time development of the cNH using the current scheme, in which no time scaling is required and so the time utilized is the “real” time [4]. The trajectory of x were within the basin of the deepest minimum at the beginning (according to the initial condition). When the temperature was increased, it escaped from the basin to approach other minima through the saddles. Note that the cNH together with the sigmoid [equation (69)] can be seen as the NH with fluctuating inverse-temperature in a range of ]*β*_L_, *β*_R_ [. In other words, as detailed in Appendix A, the cNH equation with *β*_R_ → *β*_L_ turns out to be the NH equation with the temperature 1/*k*_B_*β*_L_. Namely, fluctuations of the cNH becomes smaller as the sigmoid width *β*_R_ − *β*_L_ is small, and the cNH tends to the conventional NH equation in the zero width limit. The trajectory of the NH with the target temperature *T*_ex_ 1/*k*_B_*β*_BG_ = 1 (Q_NH_ = 0.5) is shown in Figure 5(b), implying a very long escape time against the potential energy barrier > 10.

**Figure 5.**
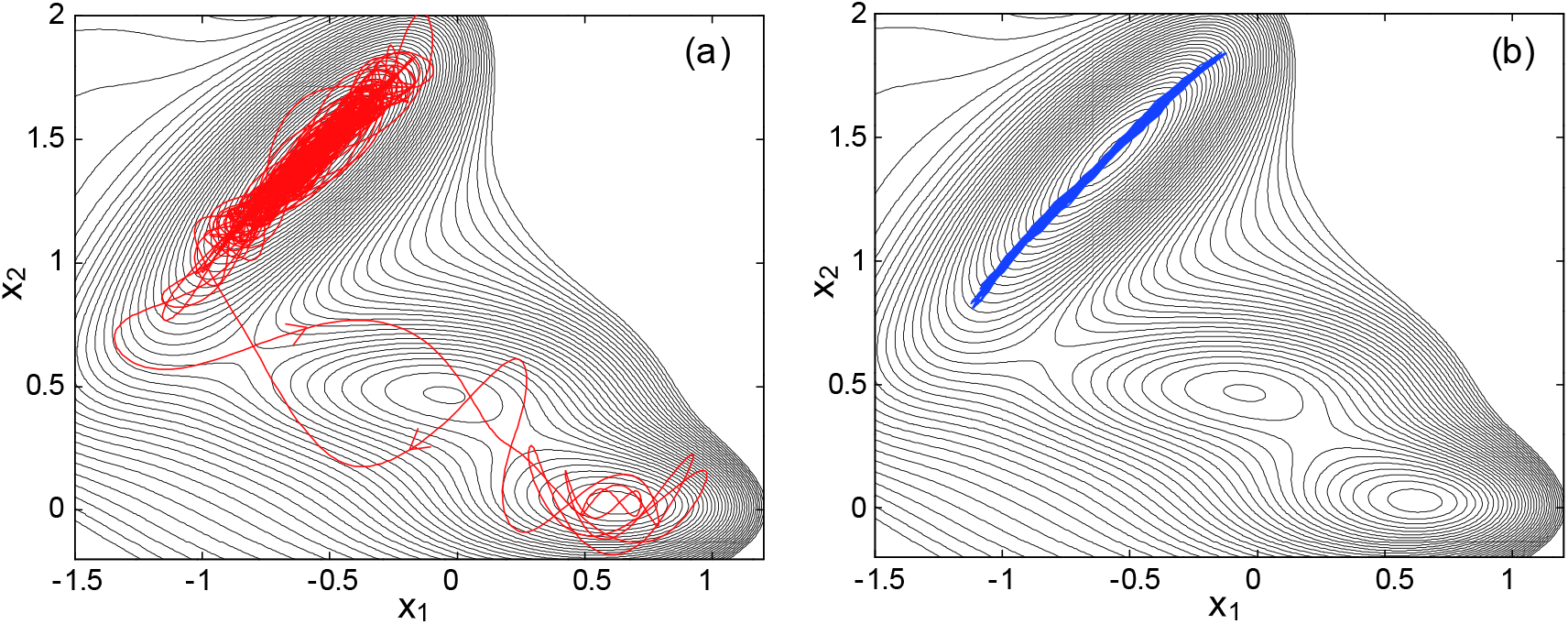
Trajectories for time 0 ≤ *t* ≤ 100 on the (*x*_1_, *x*_2_) plane for the 2D model. (a) cNH and (b) NH with temperature *T*_ex_ = 1. The initial point is *x*_1_(0) = −0.5, *x*_2_(0) = 1.5.

## VI. CONCLUSION

In the present study, we introduced a new cNH equation so as to avoid the scaled-time average in the original scheme. This was completed by a straightforward approach to obtain specific function forms so as to satisfy the Liouville equation of the total system. The success is due to the good structure of the vector field in the original cNH equation and so that of the original NH equation. Hence, we are now able to provide a fluctuating temperature environment against any physical system and also observe the realistic time development of such a physical system. It also enables the statistical description of the physical system via the invariant measure of the total system obeying the Liouville equation. We also found that the original cNH scheme with the function setting in our previous works corresponds to the current cNH scheme with modified function parameter values. The numerical investigations using the two model systems, 1D harmonic oscillator and 2D Müller-Brown potential system, showed that the correct distribution functions can be produced in the current scheme. The differences between the current and original cNH schemes in distributions and dynamics were seen in the 1HO system, as theoretically formulated. We also studied the sampling efficiency of the current method in the 2MB system via changing the parameters of the sigmoid function currently adopted to define the dynamical temperature. We compared physically meaningful trajectories obtained by the current method with those obtained by the constant temperature NH method. From these numerical results, we expect the sampling ability for realistic complex physicochemical systems and the utility for a fluctuating temperature in investigating transition states, chemical reactions, and diffusion process.

## Acknowledgments

We acknowledge the support by the “Development of core technologies for innovative drug development based upon IT” from Japan Agency for Medical Research and development, AMED. K M was supported by a Grant-in-Aid for Young Scientists (15K18520) from JSPS.

## Appendix A: features of *f* and *σ*

We first observe expectation values and relevant quantities when we use *f* defined by equation (71). In this case, for *h*: *I*_B_ → ℝ, we have

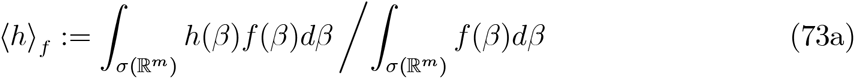

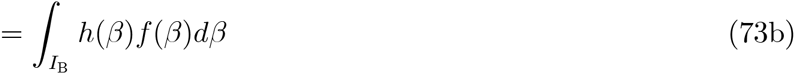

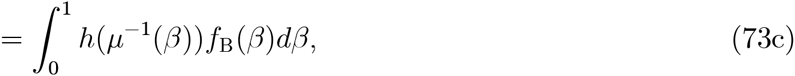

assuming the finiteness of equation (73c). Thus, for *ρ*_E_ defined by equation (61), we have

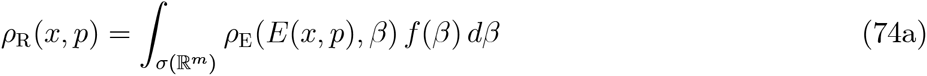

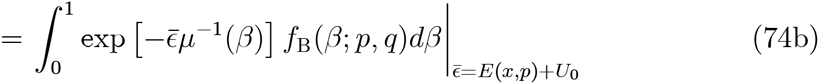

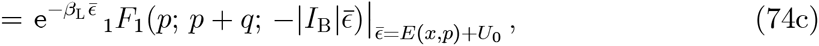

where _1_*F*_1_ is the confluent hypergeometric function, with *p* and *q* being the parameters for *f*_B_. Note that this new *f* allows any value of 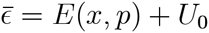 to establish *ρ*_R_(*x,p*), while *f* defined by equation (65) imposes the condition that 1 + (1/*α*_2_)(*E*(*x,p*) + *U*_0_) > 0.

We second find the explicit forms of the principal-part density *ρ*_1_ (*x,p,𝒬*), the TS potential energy *V*_*E*(*x,p*)_(*𝒬*), and the TS force −∇*V*_*E*(*x,p*)_(*𝒬*) with using *f*, *σ*, and *ρ*_E_ that are defined by equations (71), (69), and (61), respectively. First, *ρ*_1_ defined by equations (11a) and (7) becomes

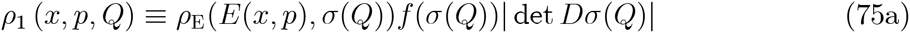

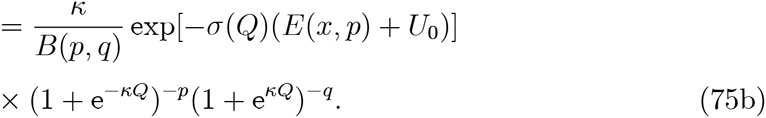

Thus we have *V*_*E*(*x, p*)_(*𝒬*) defined by equation (47) and the force − *DV*_*E*(*x,p*)_(*𝒬*) ∈ ℝ are

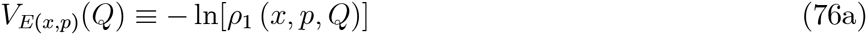

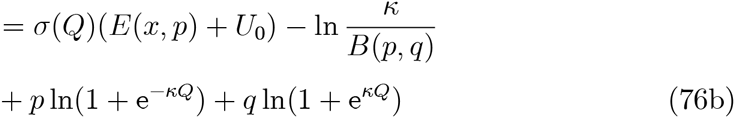

and

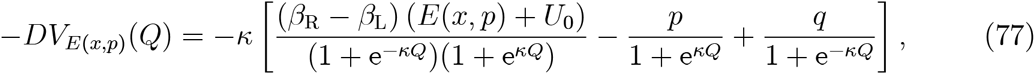

respectively. Note that new *V*_*E*(*x, p*)_(*𝒬*) defined by equation (76) is an asymptotically linear confining potential, i.e.,

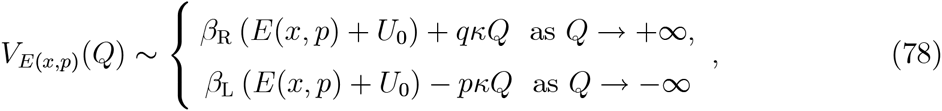

for any *p, q, κ* > 0, *β*_R_ > *β*_L_ ≥ 0, and any *E*(*x,p*) + *U*_0_ ∈ ℝ. See Figure A1.

**Figure A1.**
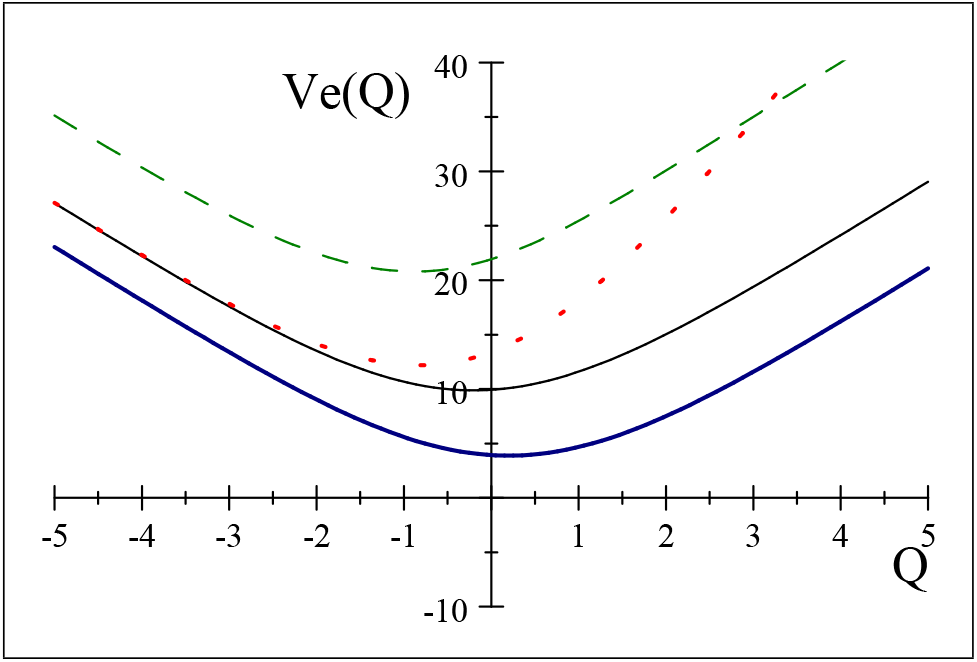
A graph of potential *V*_*E*(*x,p*)_(*𝒬*) represented by Eq. (76) (dropped ln 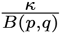) with (black solid) *p* = *q* = 5, *κ* = *β*_L_ =1, *β*_R_ =2, and *e* = *E*(*x,p*) + *U*_0_ = 2; (red dot) *q* = 10; (green dash) *e* = 10; (blue thick-solid) e = −2.

We finally discuss a physical meaning of the range of *σ* defined by equation (69), where we shall assume *β*_L_ > 0, and consider a relationship between the NH and cNH equations. Note *β*_R_ = *β*_L_ in equation (69) is forbidden, since the yielding result, *σ* = *β*_L_ =constant, contradicts Condition 1. However, if we consider a case near the limit *β*_R_ → *β*_L_, it is natural to think that cNH turns out to be the single NH with the temperature 1/*k*_B_*β*_L_, since the cNH is the NH with fluctuating inverse-temperature in a range ]*β*_L_, *β*_R_ [. In fact, in a special condition, it is true and the cNH equation is made from two decoupled NH equations. Such a condition is to use *β*_E_ defined by equation (61), *β*_Z_ defined by equation (9a), and *c_T_* = 0. Then we have *T*(*x,p,𝒬*) → 1/*k*_B_*β*_L_ = *T*_ex_ [see equation (63)], *T*(*x,p,𝒬*) *τ*_Z_ (*ζ*) → (2*c*_Z_/*β*_L_)*ζ* =const.×*ζ* [see equation (5a)], and *V_ϵ_*(*𝒬*) will be independent of *𝒬* [see equations (47) and (7), where *ρ_f_*(*𝒬*) ~ 0]. Thus, the PS approaches the NH system with the constant temperature *T*_ex_, the TS approaches the NH system of the ideal gas, and the PS and TS become decoupled.

Thus, in these function setting, the cNH with *β*_R_ → *β*_L_ is the NH with *β*_L_ (viz., the NH with temperature 1/*k*_B_*β*_L_), and similarly, the cNH with *β*_L_ → *β*_R_ is the NH with *β*_R_. Namely, the cNH gives a connection of the NH with *β*_L_ and the NH with *β*_R_. This can be translated to a statement that the cNH vector field provides a *homotopy* connecting two NH vector fields, 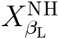 and 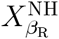, where 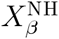 indicates the vector field of ODE (1) with *T*_ex_ = 1/*k*_B_*β* and Q_NH_ ≡ *β*/2*c*_Z_. To see this, we define

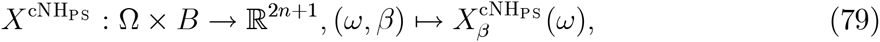

where 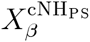 is the first three component of the vector field of equation (23), viz.,

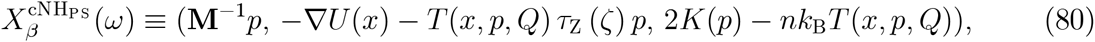

with the dynamical temperature

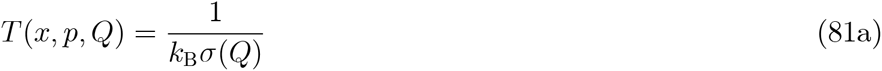

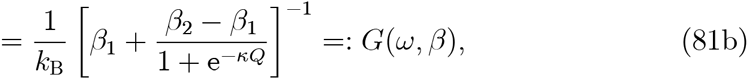

whose range is ]*β*_1_, *β*_2_[. Here we denote the both-ends inverse temperatures as *β* ≡ (*β*_1_, *β*_2_) ∈ *B*:= {(*β*_1_, *β*_2_) ∈ ℝ^2^| *β*_L_ ≤ *β*_1_ ≤ *β*_2_ ≤ *β*_R_} [Note that although the condition *β*_L_ < *β*_R_ is needed to well define the total vector field of the cNH equation (23), it is not needed to only define equation (80). Thus the definition of *B* makes sense]. Then we see that equation (79) is continuous due to the continuity of *G*. Thus, for any continuous map (path) *φ*: *I* ≡ [0, 1] → *B* such that *φ*(0) = (*β*_L_, *β*_L_) and *φ*(1) = (*β*_R_, *β*_R_), we have a continuous map,

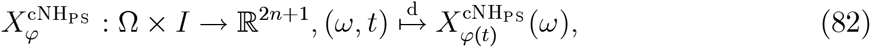

which can be identified as the homotopy from 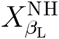 to 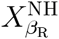, since 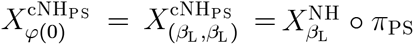 and 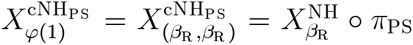, where *π*_PS_: Ω → ℝ^2*n*+1^: *ω* ↦ (*x,p, ζ*) is the projection. This is a nontrivial homotopy in the sense that it gives non NH equations except at the both ends *t* = 0 and 1. A schematic view is shown in Figure A2.

**Figure A2.**
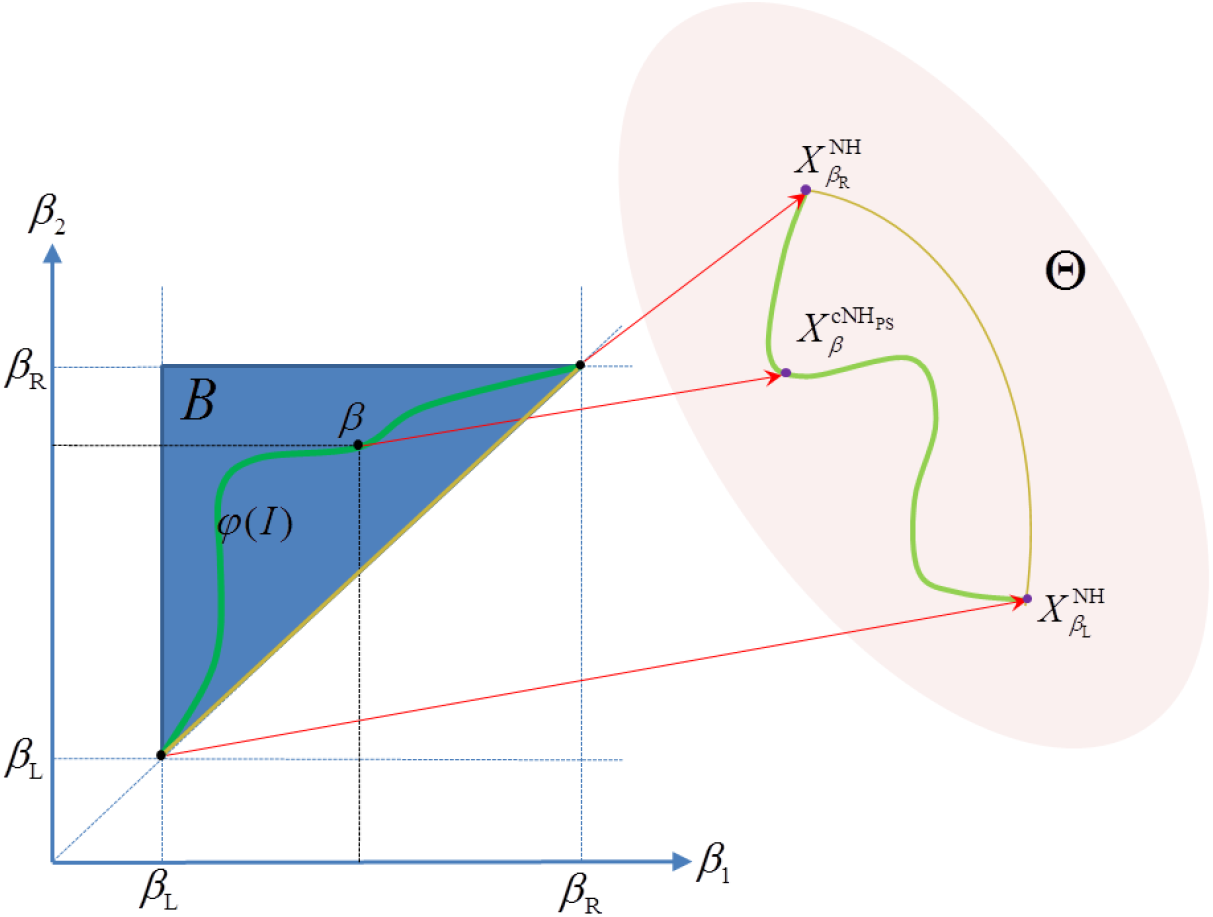
A schematic figure to represent the homotopy between the NH fields 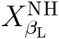 and 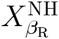 *B* is the accessible area of *β* ≡ = (*β*_1_, *β*_2_) ∈ ℝ^2^, where *β*_1_ and *β*_2_ is the infimum and supremum, respectively, of the inverse temperature of the cNH coupled with sigmoid 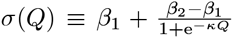. Θ represents a set of maps from Ω to ℝ^2*n*+1^. The red arrows show the mapping 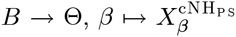. Here, 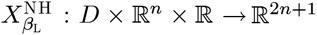 is identified with 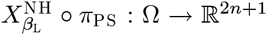, and similar for 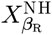. The diagonal line in *B* is mapped into the trivial (pure NH) homotopy indicated by the arc in Θ.

## Appendix B

For ODE (23), 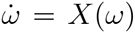, the extended space formalism [24, 25] defines an extended space Ω′ = Ω × ℝ and provides an extended ODE on it by

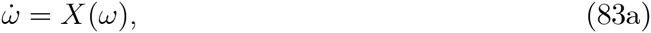

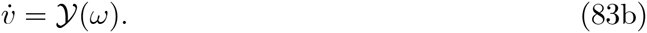

Here, *v* is introduced as an extended variable on ℝ, and

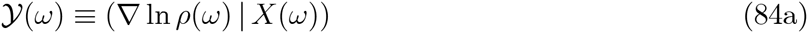

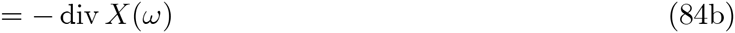

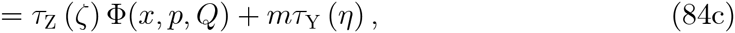

with

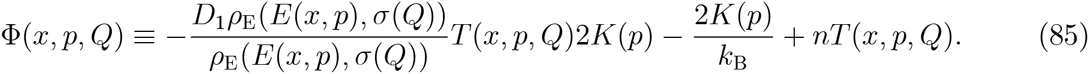

Note that Φ(*x,p, 𝒬*) = *nT*(*x,p,𝒬*) = *n*/*k*_B_*σ*(*𝒬*) if we use equation (61) and *c_T_* = 0. The ODE (83) has an invariant function [24] defined by

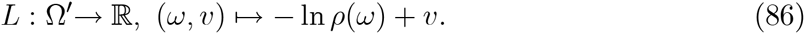

The error of the numerical integration can be checked by monitoring the value of equation (86) in the integration process.

